# Primordial germ cell DNA demethylation and development require DNA translesion synthesis

**DOI:** 10.1101/2023.07.05.547775

**Authors:** Pranay Shah, Ross Hill, Stephen Clark, Camille Dion, Abdulkadir Abakir, Mark Arends, Harry Leitch, Wolf Reik, Gerry Crossan

## Abstract

Mutations in DNA damage response (DDR) factors are associated with human infertility, which affects up to 15% of the population. It remains unclear if the role of DDR is solely in meiosis. One pathway implicated in human fertility is DNA translesion synthesis (TLS), which allows replication impediments to be bypassed. We find that TLS is essential for pre-meiotic germ cell development in the embryo. Loss of the central TLS component, REV1, significantly inhibits the induction of human PGC-like cells (hPGCLCs). This is recapitulated in mice, where deficiencies in TLS initiation (*Rev1^-/-^* or *Pcna^K164R/K164R^*) or extension (*Rev7^-/-^*) result in a >150-fold reduction in the number of primordial germ cells (PGCs) and complete sterility. In contrast, the absence of TLS does not impact the growth, function, or homeostasis of somatic tissues. Surprisingly, we find a complete failure in both activation of the germ cell transcriptional program and in DNA demethylation, a critical step in germline epigenetic reprogramming. Our findings show that for normal fertility, DNA repair is required not only for meiotic recombination but for progression through the earliest stages of germ cell development in mammals.

## Introduction

The development of germ cells and their differentiation into gametes is crucial for the faithful transmission of both genetic and epigenetic information to the next generation. Primordial germ cells (PGCs) are the first germ cells to emerge in the post-implantation embryo. In mice, as few as 3-5 founder PGCs are specified at E6.0-6.5 and these undergo extensive proliferation, increasing to around 20,000 by E12.5 (*1–3*). To develop into functional gametes, PGCs must undergo a unique developmental program, repressing somatic genes and activating pluripotency and germ-cell-specific factors (*4, 5*). This entails extensive epigenetic reprogramming resulting in altered histone modifications and DNA demethylation which facilitates the erasure of genomic imprints, and reactivation of the inactive X-chromosome (*6, 7*). DNA demethylation has been proposed to occur by multiple mechanisms (*8*). A prevalent model describes a two-step process involving a passive demethylation phase in which DNA methylation is diluted by DNA replication (‘reprogramming step 1’) followed by active DNA demethylation that occurs upon colonization of the embryonic gonads (‘reprogramming step 2’) (*9–14*). Thus, PGC development is highly dependent on DNA replication, both for lineage expansion and for epigenetic reprogramming.

The replication of DNA can be hindered by various obstacles, such as chemical damage to the DNA molecule or DNA secondary structures (*15*). The failure to resolve these impediments can have catastrophic consequences for a cell. Incomplete or under-replication of the genome can block cell cycle progression, either directly or by activation of the DNA damage response (DDR) and cell cycle checkpoints (*16*). If this persists the cell may ultimately die. To combat these challenges, eukaryotes have evolved both DNA repair as well DNA damage tolerance (DDT) mechanisms. One route of DDT is error-prone translesion synthesis (TLS). TLS allows DNA replication to continue past impediments and facilitates the filling of gaps which remain at the end of S-phase (*17*). TLS utilizes specialized polymerases with active sites that can accommodate damaged or distorted DNA templates and which lack proofreading activity, hence increasing the risk of DNA mutagenesis (*18, 19*). In the germ cell compartment, ensuring that replication can proceed is of paramount importance to allow sufficient cellular expansion to guard against sterility. However, any increase in the mutagenicity of replication increases the risk of deleterious phenotypes and inherited disease in future generations.

Recent genome wide association studies (GWAS) have implicated multiple DDR pathways as determinants of infertility in humans, however little is known about the underlying mechanism (*20–22*). In this study, we investigate one of those pathways, TLS, and find it plays a crucial role in the development of embryonic germ cells in both humans and mice. Our results show that factors involved in sequential stages of TLS are essential for PGC development. In the absence of TLS, PGCs are specified in normal numbers but fail to expand, resulting in a >150-fold reduction. In contrast to the severe effect on the germline, somatic tissues of these mutants are unperturbed in their development with no discernible impact on homeostasis or survival. Consistent with a defect in TLS, mutant PGCs show reduced proliferation, accumulation at the G2/M phase of the cell cycle and increased markers of unresolved DNA damage. In addition, the loss of TLS prevents progression of the germ cell transcriptional program and results in failure of genome-wide DNA demethylation, an essential and conserved step in the development of mammalian embryonic germ cells. Our findings define a critical role of TLS specifically in germ cell development, safeguarding fertility and enabling successful PGC epigenetic reprogramming.

## Results

### REV1 is required for PGC development in human and mouse

GWAS studies focused on infertility have revealed genes important for successful human germline development (*20–22*). Notably, factors that deal with DNA damage are frequent hits. The DDR has been implicated in germ cell production and maintenance, including Fanconi Anemia crosslink repair, base excision repair and homologous recombination (*23–25*). While TLS factors have been implicated as determinants of human fertility however, why they are needed has not been explained. TLS is important for the completion of DNA replication, and we therefore hypothesized that it may play a role in ensuring successful genome duplication during highly proliferative stages of gametogenesis. As early embryonic germ cell development is particularly proliferative, we asked if TLS is required during primordial germ cell (PGC) development. We employed an *in vitro* model in which iPSCs are differentiated into human PGC-like cells (hPGCLCs) (**Fig. 1a**) (*26*). To study the potential roles in human germ cell development we focused on REV1, a core component of TLS (*27–30*). REV1 was disrupted in human BTAG iPSCs carrying both the BLIMP1-tdTomato and TFAP2C-EGFP PGCLC reporters (**Supplementary Fig. 1a**) (*26*). Clones with successful disruption of REV1 were identified by PCR then validated by testing for hypersensitivity to mitomycin C (MMC) (**Supplementary Fig. 1b-c**). Consistent with a role in embryonic germ cell development, we found that three independent *REV1^-/-^* lines had a significant reduction in the ability to induce hPGCLCs (**Fig. 1b-c and Supplementary Fig. 1d**). While the frequency of hPGCLCs generated was significantly reduced in the absence of REV1, the few induced hPGCLCs expressed canonical germ cell factors (**Fig. 1d**).

**Fig. 1.**
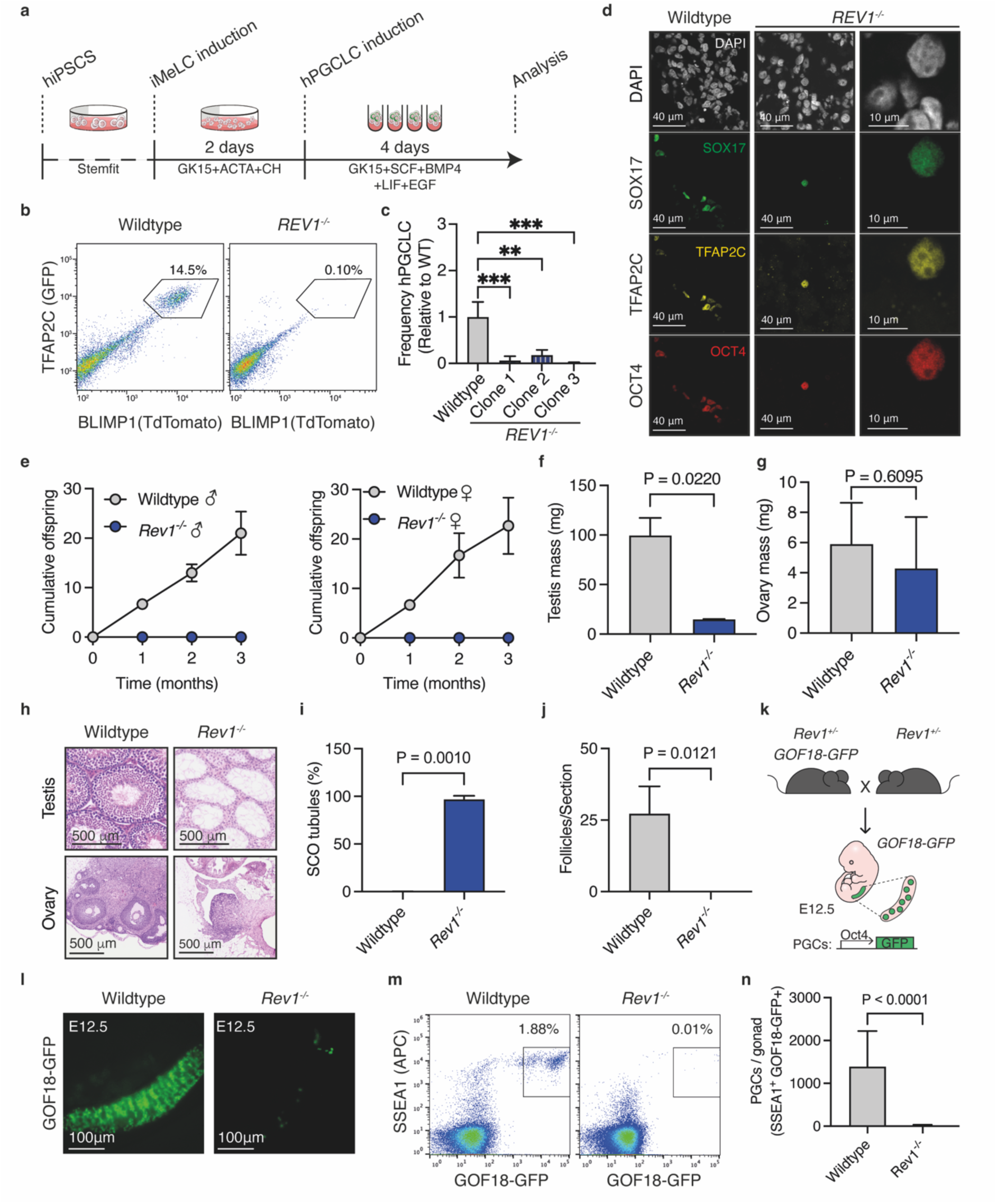
REV1-deficiency leads to defects in hPGCLC induction and infertility in mice due to compromised PGC development. **(a)** Schematic of the hPGCLC differentiation adapted from (*26*) **(b)** Representative flow cytometry plots of hPGCLC percentages at day 4 of aggregate differentiation from wildtype or *REV1^-/-^* BTAG cells. Box shows the percentage of TFAP2C-GFP and BLIMP1-tdTomato double positive cells. **(c)** Quantification of hPGCLC (TFAP2C-GFP^+^ BLIMP1-tdTomato^+^) frequency from three independent wildtype or *REV1^-/-^* clones. *REV1^-/-^* clones were each differentiated three times. Each experiment was normalized to wildtype (Anova test, **:P ≤ 0.01 and ***: P ≤ 0.001) **(d)** Representative images of wildtype and *REV1^-/-^* aggregate immunostained for OCT4, SOX17 and TFAP2C. **(e)** Cumulative number of offspring when male (left) and female (right) wildtype or *Rev1^-/-^* mice were mated with wildtype mates of the opposite sex (n=6 mice per genotype, 3 per sex). **(f)** Quantification of testis mass of 8-12-week-old wildtype and *Rev1^-/-^* mice (n=12, 2, left to right). **(g)** Quantification of ovary mass from 8-12-week-old wildtype and *Rev1^-/-^* mice (n=13, 2, left to right). **(h)** H&E-stained ovaries and testes seminiferous tubules from 8–12-week-old wildtype and *Rev1^-/-^* mice (similar results were obtained from 3 independent animals per genotype and sex). **(i)** Quantification of SCO tubules per testis of 8-12-week-old wildtype and *Rev1^-/-^* mice (n=10 and 4 independent animals, left to right). **(j)** Quantification of follicles per section of ovary from 8-12-week-old wildtype and *Rev1^-/-^* mice (n=8 and 3 independent animals, left to right). **(k)** Schematic for generation of *Rev1^-/-^* embryos harbouring the GOF18-GFP reporter. **(l)** GFP fluorescence images of gonads from wildtype and *Rev1^-/-^* E12.5 embryos. **(m)** Representative flow cytometry plots of PGCs (SSEA1^+^GOF8-GFP^+^) from wildtype and *Rev1^-/-^* E12.5 embryos. **(n)** Quantification of PGCs by flow cytometry from wildtype and *Rev1^-/-^* E12.5 embryos (n=35 and 12, left to right).

Due to limitations of studying human gametogenesis *in vivo*, we asked if the requirement for TLS factors in PGC development was conserved to mice, thus facilitating mechanistic studies. REV1-deficient mice were crossed with wildtype mates to assess fertility. Consistent with our observations in human, we found that neither male nor female *Rev1^-/-^* mice gave rise to offspring despite evidence of copulation (**Fig. 1e**) (*31*). Upon analysis of the gonads, we observed a striking reduction in testis mass (**Fig. 1f-g**). Histological analysis of the testes revealed a majority (96.8%) of Sertoli-cell-only (SCO) seminiferous tubules and a complete absence of promyelocytic leukemia zinc finger (PLZF)^+^ cells, which marks undifferentiated spermatogonial stem cells (SSCs, **Fig. 1h-i and Supplementary Fig. 1f**) (*32*). In females, we observed no follicles upon histological examination of *Rev1^-/-^*ovaries, suggesting a complete failure of oogenesis (**Fig. 1h** and **j**). Together, these data reveal a lack of gametes in both *Rev1^-/-^* male and female mice. The lack of visible PLZF^+^ cells in male gonads and meiotic cells in either sex argues that the defect is pre-meiotic which is consistent with the hPGCLC data in suggesting a failure during embryonic germ cell development. Therefore, we studied the development of embryonic germ cells using the GOF18-GFP PGC reporter in which GFP expression is driven by a fragment of the *Oct4* promoter (Oct4ΔPE) (**Fig. 1k**) (*33, 34*). We intercrossed *Rev1^+/-^* mice carrying the GOF18-GFP PGC reporter and harvested the embryonic gonads at E12.5, prior to sexually divergent germline development. Imaging of the gonads revealed a dramatic reduction in GOF18-GFP^+^ cells in *Rev1^-/-^* embryos compared to wildtype controls, consistent with the hPGCLC data (**Fig. 1l**). Quantification of PGCs (SSEA1^+^GOF18-GFP^+^) by flow cytometry revealed a significant reduction in *Rev1^-/-^* embryos (**Fig. 1m-n**). These data show that the TLS factor REV1 plays a key role in embryonic germ cell development in both humans and mice.

### The requirement for the C-terminal domain of REV1 and REV7 links TLS to PGC development

REV1 can act enzymatically through its deoxycytidyl transferase activity which can drive mutagenesis and is important for immunoglobulin diversification (*27, 29, 30*). Alternatively, it can act as a protein-scaffold through its C-terminal protein interaction domain which is able to recruit TLS factors to sites of lesion bypass (*35*). This region of REV1 plays a crucial role in maintaining cellular resistance to DNA damage that impedes replication indicating that the C-terminus is the key region of REV1 for TLS transactions. To determine if the catalytic or recruitment function of REV1 is required for PGC development, we generated a catalytically dead allele of *Rev1*, in which the catalytic residues D568 and E569 were mutated to alanine (*Rev1^AA^*) (**Fig. 2a** and **Supplementary Fig. 2a**) (*36*). We also generated mice with a mutant *Rev1* allele in which the DNA encoding the final 100 amino acids of REV1 was deleted (*Rev1^CT^*) (**Fig. 2a** and **Supplementary Fig. 2b-c**). To validate these *Rev1* mutant alleles, we derived cell lines from mice and measured mRNA expression and cellular sensitivity to DNA damaging agents. The mRNA expression of mutant alleles was comparable to that of *Rev1* in wildtype cells (**Supplementary Fig. 2d**). Moreover, the C-terminus but not catalytic activity of REV1 was required to overcome replication blocking DNA damage, in line with previous reports (**Supplementary Fig. 2e**).

**Fig. 2.**
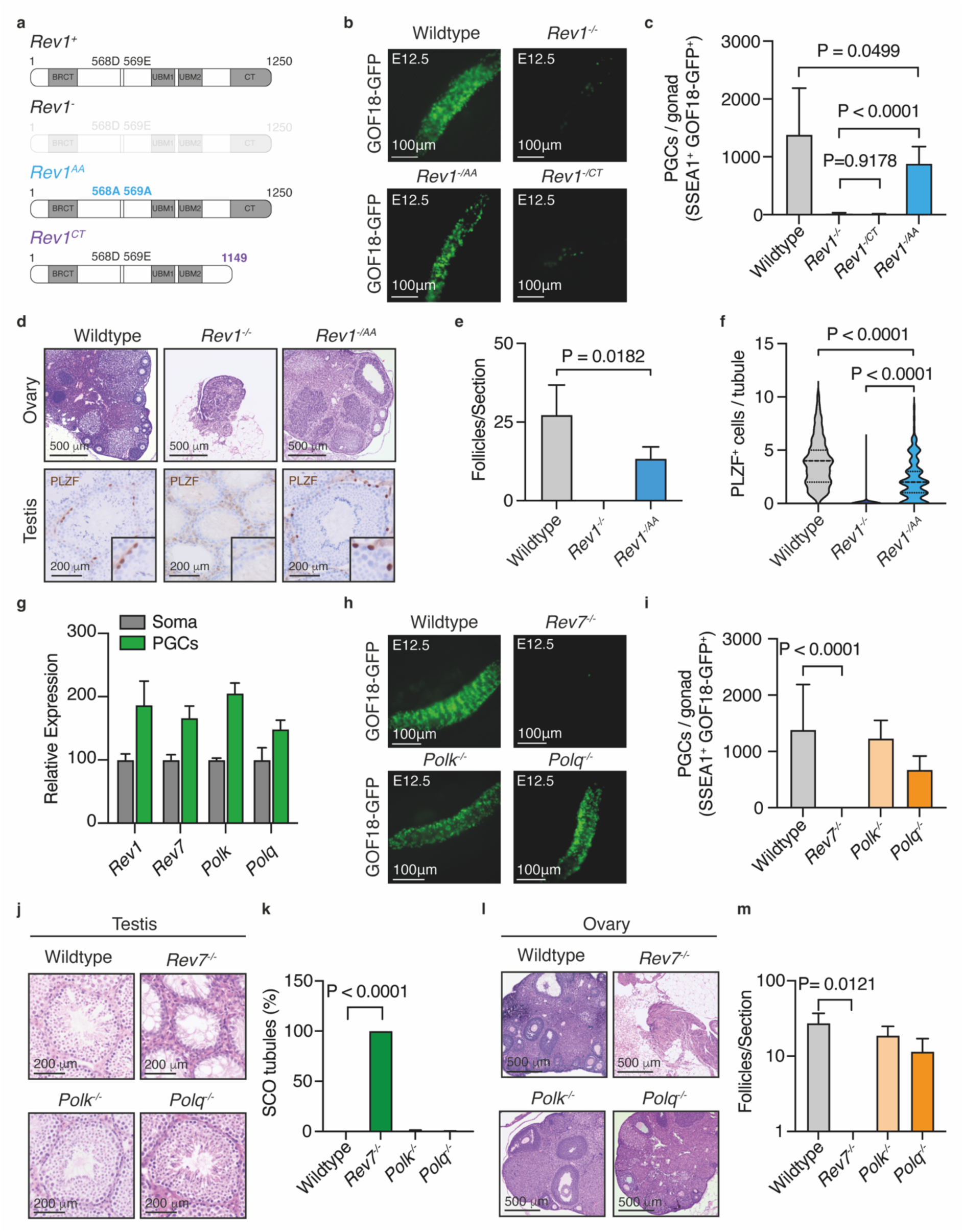
REV7 is required for PGC development in mice. **(a)** Schematic of wildtype (*Rev1^+^*), null (*Rev1*^-^), catalytically inactive REV1 (*Rev1^AA^*) and C-terminally truncated REV1 (*Rev1^CT^*) alleles. **(b)** GFP fluorescence images of gonads from wildtype, *Rev1^-/-^*, *Rev1^-/AA^*and *Rev1^-/CT^* E12.5 embryos. **(c)** Quantification of PGCs by flow cytometry from wildtype, *Rev1^-/-^*, *Rev1^-/AA^* and *Rev1^-/CT^* embryos at E12.5 (n=35, 12, 10 and 7, left to right). **(d)** H&E-stained ovaries and PLZF-stained testes from 8–12-week-old wildtype, *Rev1^-/-^* and *Rev1^-/AA^* mice (similar results were obtained from 3 independent animals per genotype and sex). **(e)** Quantification of follicles per section of ovary from 8-12-week-old wildtype, *Rev1^-/-^* and *Rev1^-/AA^* mice (n=8, 3 and 3 independent animals, left to right). **(f)** Frequency of PLZF^+^ cells per seminiferous tubule of 8-12-week-old wildtype, *Rev1^-/-^*and *Rev1^-/AA^* mice (the data shown represent the median and interquartile range; n=150 tubules per genotype, 50 per genotype). **(g)** Droplet digital PCR (ddPCR) gene expression analysis *Rev1*, *Rev7, Polk* and *Polq* in FACS-purified PGCs and surrounding somatic cells (SSEA1^-^GOF18-GFP^−^) from E10.5 embryos (n=3 independent embryos). **(h)** GFP fluorescence images of gonads from wildtype, *Rev7^-/-^*, *Polk^-/-^*and *Polq^-/-^* E12.5 embryos. **(i)** Quantification of PGCs from wildtype, *Rev7^-/-^*, *Polk^-/-^* and *Polq^-/-^* E12.5 embryos by flow cytometry (n=35, 14, 8 and 9, left to right). **(j)** H&E-stained testis seminiferous tubules from 8–12-week-old wildtype and mutant mice. **(k)** Quantification of SCO tubules per testis of 8-12-week-old mice (n=10, 7, 8 and 14, left to right). **(l)** H&E-stained ovaries from 8–12-week-old wildtype and mutant mice. **(m)** Quantification of follicles per section of ovary from 8-12-week-old mice (n=8, 3, 4 and 11, left to right).

We crossed the *Rev1^AA^* or *Rev1^CT^* alleles with *Rev1^+/-^* mice carrying the GOF18-GFP PGC reporter. At E12.5, we found that *Rev1^-/AA^* embryos had a comparable number of PGCs to wildtype (median wildtype = 1383, median *Rev1^-/AA^* = 884, **Fig. 2b-c**). These PGCs generated mature gametes in the gonads of adult *Rev1^-/AA^* mice which were competent in giving rise to viable offspring (**Fig. 2d-f and Supplementary Fig. 2f**). In contrast, E12.5 *Rev1^-/CT^* embryos showed a significant reduction in the number of PGCs, with similar numbers to those in *Rev1^-/-^* embryos (median = 27 and 20, respectively, **Fig. 2b-c**). Together these data show that whilst the catalytic activity of REV1 is dispensable for fertility, the C-terminus, which coordinates protein-protein interactions during TLS, is critical for PGC development.

To investigate the function of TLS in PGC development further, we investigated known REV1 interactors. First, we measured the expression of a subset of interactors (REV7, POLΚ) and POLΘ and found that like *Rev1*, each was more highly expressed in PGCs than in the surrounding somatic cells (**Fig. 2g**) (*29, 37*). To determine if these factors are also required for PGC development, we generated E12.5 embryos deficient in each factor and analyzed them using the GOF18-GFP PGC reporter (**Fig. 2h-i**). Whilst *Polk^-/-^* and *Polq^-/-^* embryos had comparable numbers of PGCs to wildtype, REV7-deficient embryos had a significant reduction, similar to that observed in *Rev1^-/-^* embryos (**Fig. 2h-i**). Consistent with this, the profound reduction in the number of PGCs in *Rev7^-/-^*embryos resulted in adult gonads devoid of germ cells (**Fig. 2j-m and Supplementary Fig. 3a**).

Together these data reveal that it is the C-terminal protein interaction domain of REV1 and its interactor REV7 that are required during PGC development. REV7/repro22 has previously been identified in an ENU screen as a factor needed for fertility and subsequent studies have shown REV7 to be important during PGC expansion (*38–40*). REV7 is the non-catalytic subunit of Polζ, a B-family polymerase that extends the nascent strand following lesion bypass before handing back to replicative polymerases (*41, 42*). As only Polζ has such an activity it is considered essential for TLS transactions. However, it also has non-TLS roles in DSB repair, cell cycle regulation and the shelterin complex (*43–46*). Crucially, Polζ is recruited to sites of lesion bypass for DNA synthesis through physical interactions between the REV7 subunit and REV1’s C-terminal domain (*47*). Hence, our discovery that both the C-terminus of REV1 and REV7 are required in PGCs argues that the common function of both - in TLS - is needed during germ cell development.

### The post-translational modification of PCNA preserves PGC development

If TLS is critical for the development of PGCs, we hypothesized that PCNA ought to play an important role. PCNA is an essential component of the replisome that links DNA repair and DDT responses to replication (*48*). Upon stalling of replication forks, lysine 164 of PCNA can be sumoylated, mono-, or poly-ubiquitinated to engage DDT pathways (*46, 48–51*). Monoubiquitination of PCNA at lysine 164 is required for TLS, facilitating the recruitment of specialized polymerases to sites of replication blocking lesions, enabling their bypass (*52–54*). As our results found a requirement for the TLS factors REV1 and REV7, we wanted to study if there is also a role for PCNA K164 modification during PGC development. We used CRISPR/Cas9 to generate mice in which lysine 164 of PCNA is mutated to arginine (*Pcna^K164R^*) (**Supplementary Fig. 4a**). In this model, PCNA K164-modification dependent DDT, including TLS, is abolished (*48*). We first set out to determine if our allele recapitulated features of previously described mice carrying the PCNA K164R mutation (*Pcna^tm1Jcbs^* MGI:3761720 and *Tg^(Pcna*K164)1Mdsc^* MGI:97503) (*55, 56*). As expected, *Pcna^K164R/K164R^* (herein referred to as *Pcna^R/R^*) mouse embryonic fibroblasts (MEFs) derived from our allele were hypersensitive to ultraviolet irradiation (*57*) (**Supplementary Fig. 4b**). However, despite PCNA K164 being important for the cellular response to replication blocking DNA damage, *Pcna^R/R^* mice were born at the expected Mendelian ratios with adult mice having similar lifespan to wildtype littermates (**Supplementary Fig. 4c-d**). Consistent with previously published data, we found that the gonads of homozygous adult mice were smaller than wildtype and that both male and female *Pcna^R/R^* mice were sterile (**Fig. 3a-d**) (*55*). Histological analysis revealed that the majority (98.2%) of the testes contained Sertoli-cell-only seminiferous tubules and that the ovaries were devoid of follicles (**Fig. 3e-h**). Similar to REV1-deficiency, the failure in gametogenesis began during embryonic development (**Fig. 3i-j**). At E12.5, the numbers of PGCs observed in *Pcna^R/R^* embryos was similar to *Rev1^-/-^*with a >150-fold reduction when compared to wildtype (**Fig. 3k**). To study if there were sex-specific differences in the embryonic germ cell defect of TLS-deficient embryos, we compared E12.5 male and female *Pcna^R/R^* embryos. We found comparable PGC numbers in both sexes, confirming a common defect (**Supplementary Fig. 4e**).

**Fig. 3.**
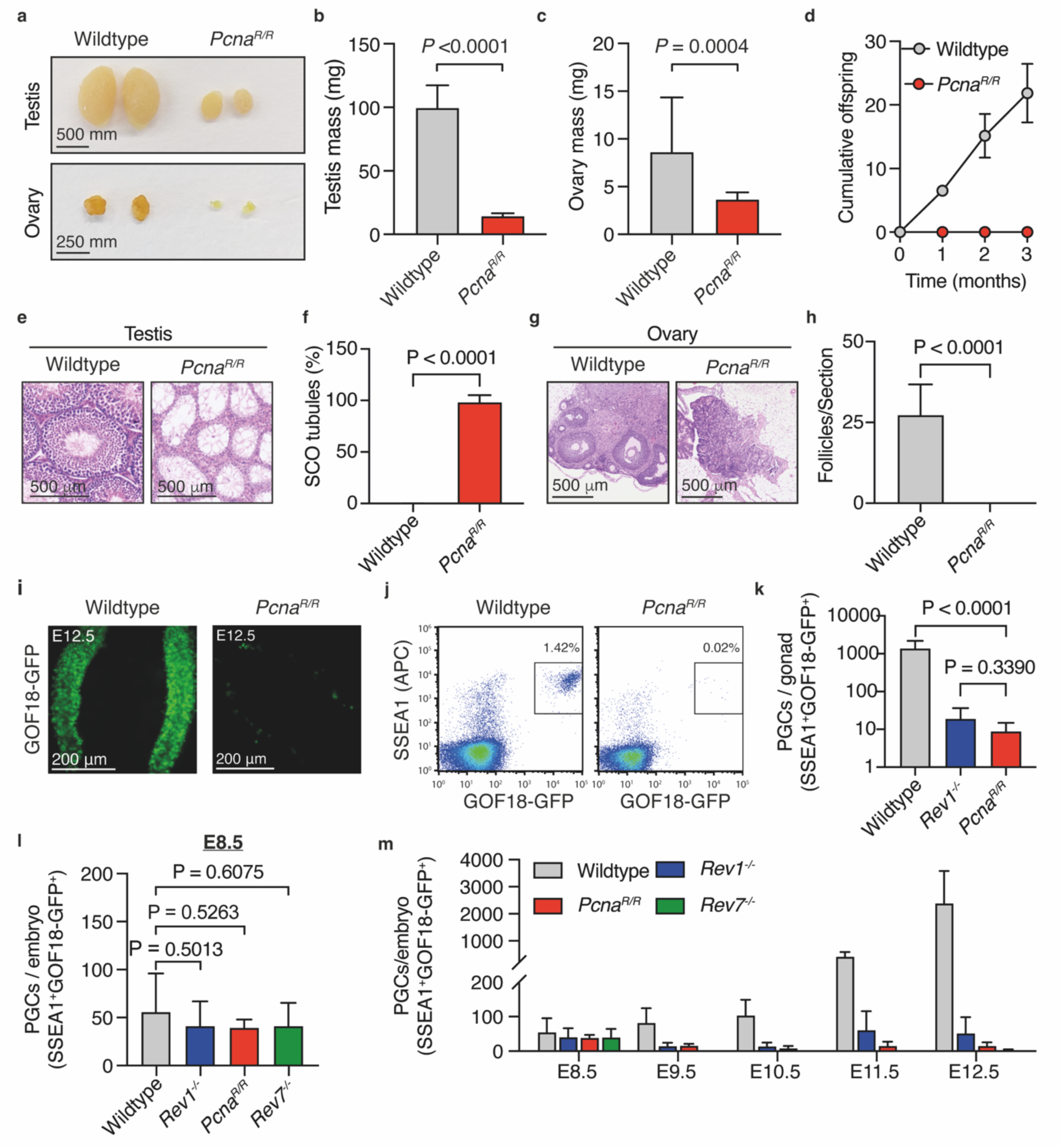
Embryonic origin of sterility upon PCNA K164 mutation. **(a)** Representative macroscopic images of testes and ovaries from wildtype and *Pcna^R/R^* mice. **(b)** Quantification of testicular mass from 8-12-week-old wildtype and *Pcna^R/R^* mice (n=12 and 10, left to right). **(c)** Quantification of ovarian mass from 8-12-week-old wildtype and *Pcna^R/R^* mice (n=13 and 8, left to right). **(d)** Cumulative number of offspring when wildtype or *Pcna^R/R^* mice were mated with wildtype mates of the opposite sex (n=6 mice per genotype, 3 per sex). **(e)** H&E-stained testis seminiferous tubules from 8–12-week-old wildtype and *Pcna^R/R^*mice (similar results were obtained from 3 independent animals per genotype and sex). **(f)** Quantification of SCO tubules per testis of 8-12-week-old wildtype and *Pcna^R/R^* (n=10 and 15, left to right). **(g)** H&E-stained ovaries and testis seminiferous tubules from 8–12-week-old wildtype and *Pcna^R/R^* mice (similar results were obtained from 3 independent animals per genotype and sex). **(h)** Quantification of follicles per section of ovary from 8-12-week-old wildtype and *Pcna^R/R^* mice (n=8 and 11, left to right). **(i)** GFP fluorescence images of gonads from wildtype and *Pcna^R/R^* E12.5 embryos. **(j)** Representative flow cytometry plots of wildtype and *Pcna^R/R^* E12.5 gonads. **(k)** Quantification of PGCs by flow cytometry from wildtype, *Rev1^-/-^* and *Pcna^R/R^* gonads at E12.5 (n=35, 12 and 21, left to right). **(l)** Quantification of PGCs by flow cytometry from wildtype, *Rev1^-/-^*, *Pcna^R/R^* and *Rev7^-/-^*embryos at E8.5 (n=18, 5, 2 and 3, left to right). **(m)** Quantification of PGCs by flow cytometry from E8.5 to E12.5 in wildtype and mutant embryos (wildtype, n = 18, 12, 9, 3 and 17; *Rev1^−/−^*, n = 5, 6, 5, 2 and 8; *Pcna^R/R^*, n = 2, 4, 4, 7 and 9; *Rev7^−/−^*, n = 3, 0, 0, 0 and 7, independent embryos, left to right).

These findings led us to investigate the timing of the PGC defect in *Rev1^-/-^*, *Rev7^-/-^* and *Pcna^R/R^* embryos more systematically. The PGCs in mouse embryos are specified between E6.0-6.5 then start to migrate and extensively proliferate from E8.5 (*1*). We first counted the number of PGCs at E8.5 and found no significant difference in the number of PGCs between wildtype and the three TLS mutants (**Fig. 3i**). We confirmed this result using an alternative PGC-reporter, *Stella-GFP* (**Supplementary Fig. 4f**) (*57*). Next, we quantified the number of PGCs by flow cytometry at each day of development between E8.5-12.5 and found that from E9.5 onwards *Rev1^-/-^* and *Pcna^R/R^*embryos had a significantly contracted PGC pool when compared to wildtype (**Fig. 3m** and **Supplementary Fig. 4g-i**). These data reveal that in addition to similar magnitudes, the timing of PGC defect is comparable across the different TLS mutants.

Having observed an essential requirement for TLS factors in mammalian germ cell development, we wondered if this was unique to the germline. In agreement with a previous report, we found a 2.6-fold reduction in the frequency of hematopoietic stem and progenitor cells (HSPCs) in adult mice (**Supplementary Fig. 5a-b**) (*58*). As adult HSCs are largely quiescent we hypothesized that the defect may begin during embryonic development when HSPCs are highly proliferative (*59*). Similar to adult mice, we found that E12.5 *Pcna^R/R^* embryos had a 1.73-fold reduction in the frequency of HSPCs, revealing that the blood stem cell defect begins during embryonic development (**Supplementary Fig. 5c**). Despite the HSPC defect, we found that blood compartment homeostasis was intact with adult *Pcna^R/R^*mice showing comparable bone marrow cellularity to littermates, normal blood cell maturation, and sustained peripheral blood homeostasis (**Supplementary Fig. 5d-g**).

We next examined a panel of vital somatic tissues and found no gross histological abnormalities consistent with the normal longevity of *Pcna^R/R^* mice (**Supplementary Fig. 6a**). As PCNA is an essential component of the replication machinery, and its modification at K164 directly couples DDT to the replisome, we assessed the proportion of actively dividing cells in highly mitotic tissues. We found comparable numbers of Ki67^+^ cells in the bone marrow, crypts of the ileum and hair follicle bulges of the skin of *Pcna^R/R^* mice compared to controls (**Supplementary Fig. 6b-d**). Furthermore, we also assessed the effect of PCNA K164R mutation on the function of the liver and kidney and found no difference compared to wildtype (**Supplementary Fig. 6e-h**). These data show that the K164R mutation of PCNA does not lead to a global reduction in proliferation nor to loss of tissue homeostasis or function. In contrast, histological analysis of the adult gonads revealed complete loss of homeostasis (**Supplementary Fig. 7**). Consistent with the histological analysis, we observe loss of tissue function in mutants leading to disruption of the hypothalamic-pituitary-gonadal axis. This led to systemic hormonal dysregulation with elevated levels of luteinising hormone (LH) and follicle stimulating hormone (FSH) driving reactive stromal hyperplasia and a significant increase in testicular interstitial cells and ovarian stroma. As animals age, persistent hypergonadotrophism leads to a substantial increase in ovarian mass with mutant ovaries of 12-month-old mice being 3 times larger than wildtype controls (**Supplementary Fig. 7d**). These features are consistent across TLS-deficient mice with similar defects seen in *Rev1^-/-^* and *Rev7^-/-^* adults (**Supplementary Fig. 7**).

Overall, whilst the PCNA K164R mutation results in increased sensitivity to exogenous DNA damage and a reduction in HSPCs, our findings reveal that the development and homeostasis of somatic tissues are largely unaffected. In contrast, the modification site of PCNA is essential for PGC development and fertility. Failure of embryonic germ cell development results in dysregulation of hypothalamic-pituitary-gonadal hormone regulation with inappropriate stromal proliferation in adults. The temporality and magnitudes of *Rev1^-/-^*, *Rev7^-/-^* and *Pcna^R/R^* germ cell defects and the molecular dissection of REV1 suggest a common origin of the defect - likely their shared role in TLS.

### Loss of genome stability and reduced proliferation of *Pcna^R/R^* and *Rev1^-/-^* PGCs

Next, we investigated the mechanisms of PGC failure in the absence of TLS factors. As TLS mitigates replication blocks, we asked if *Pcna^R/R^* or *Rev1^-/-^*PGCs accumulate unresolved DNA damage. We stained PGCs for the phosphorylation of histone variant H2A.X (ɣ-H2A.X), a marker of DNA double-strand breaks (*60*). A greater proportion of TLS-deficient PGCs had >10 ɣ-H2A.X foci when compared to wildtype (**Fig. 4a**). Consistent with our previous data this finding was specific to PGCs as an increase in foci was not observed in surrounding somatic tissue (**Supplementary Fig. 8a**). To build on this, we stained for RPA, which binds to single stranded DNA generated during repair transactions or when DNA replication is perturbed and leads to under-replicated regions in the genome (*61*). We observed a higher frequency of PGCs with nuclear RPA foci in *Pcna^R/R^* and *Rev1^-/-^* embryos (**Fig. 4b**). This shows that PGC development requires TLS and that in its absence cells accumulate damaged DNA.

**Fig. 4.**
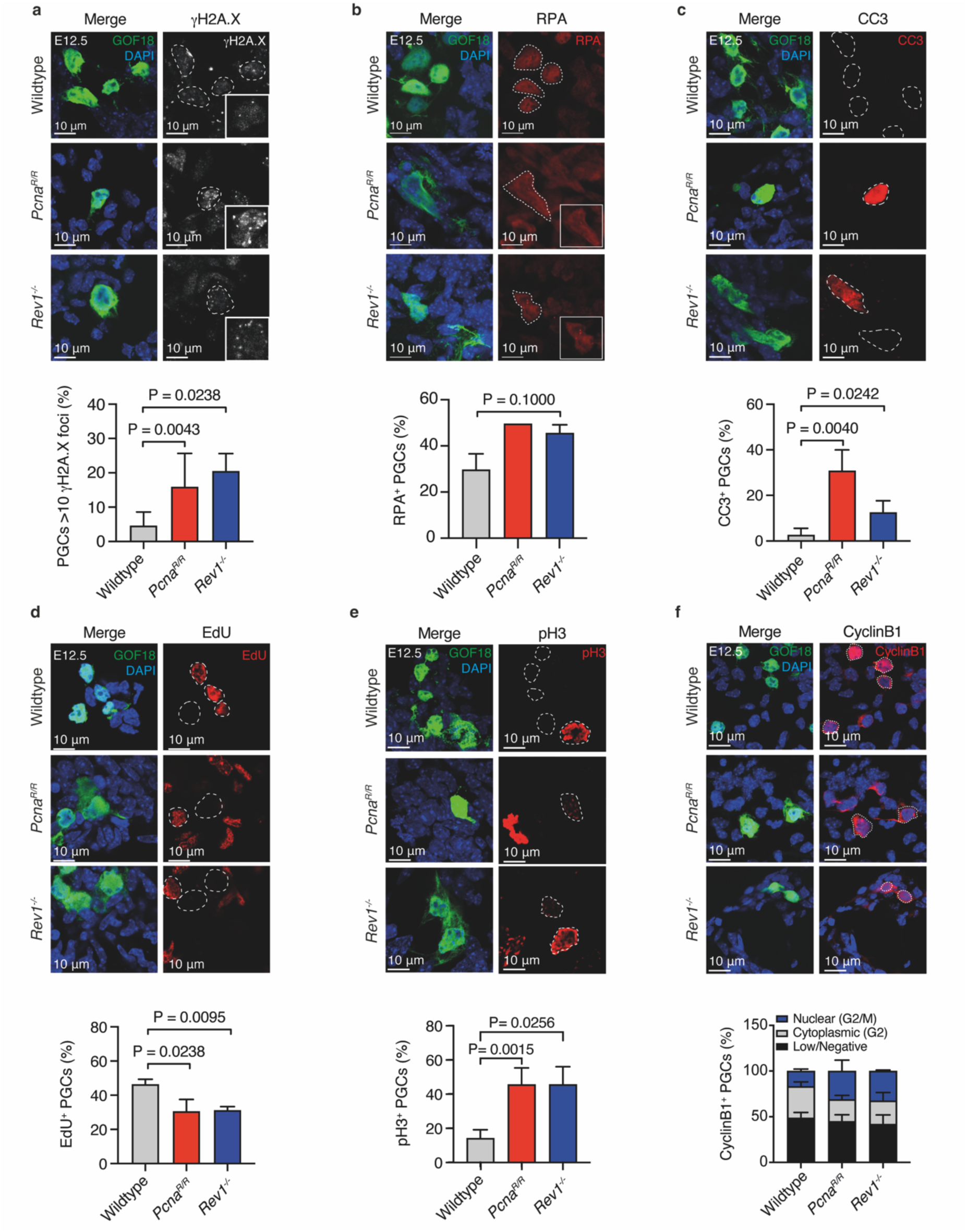
Genome instability and cell cycle abnormalities in *Pcna^R/R^* and *Rev1^-/-^* PGCs. **(a)** Top: Representative images of ɣ-H2A.X foci in the nucleus of PGCs (GOF18-GFP^+^) at E12.5. Bottom: Frequency of PGCs with >10 ɣ-H2A.X foci per nucleus (n=6, 5 and 3, left to right). **(b)** Top: Representative images of RPA foci in the nucleus of PGCs at E12.5. Bottom: Frequency of PGCs with RPA foci (n=3, 1 and 3, left to right). **(c)** Top: Representative images of E12.5 gonads stained for cleaved-Caspase-3 (CC3) and GFP. Bottom: Frequency of PGCs that stain positive for CC3 (n=8, 4 and 3, left to right). **(d)** Top: Representative images of E12.5 gonads stained for EdU and GFP . Bottom: Frequency of PGCs that stain positive for EdU (n=3 per genotype). **(e)** Top: Representative images of E12.5 gonads stained for phosphorylated-histone-H3 (pH3) and GFP. Bottom: Frequency of PGCs that stain positive for pH3 (n=11, 4 and 2, left to right). **(f)** Top: Representative images of E12.5 gonads stained for cyclin B1 and GFP. Bottom: Frequency of PGCs with nuclear, cytoplasmic or low/negative cyclin B1 staining (n=5, 3 and 3 individual embryos, left to right).

Altered proliferation and apoptosis are frequent cellular responses to persistent DNA damage. The failure of the PGC pool to expand between E8.5 - E12.5 in TLS-deficient embryos could be explained by either reduced PGC proliferation, increased PGC death, or a combination of both. We therefore first asked if TLS-deficiency leads to increased PGC death by staining E12.5 urogenital ridges for the apoptotic marker cleaved-caspase 3 (CC3) (*62*). We observed significant 7.7-fold and 5.1-fold increases in the proportion of CC3-positive PGCs in *Pcna^R/R^* and *Rev1^-/-^* embryos respectively (**Fig. 4c**). In contrast, there was no significant increase in the proportion of apoptotic (CC3^+^) somatic cells in either mutant (**Supplementary Fig. 8B**). We next assessed *in vivo* PGC proliferation by injecting pregnant dams with a single dose of ethynyl-2’-deoxyuridine (EdU), which is incorporated into the DNA of replicating cells allowing the quantification of the proportion of cells in S-phase during the EdU pulse (**Supplementary Fig. 8c**) (*63*). We found a significant reduction in the frequency of EdU^+^ PGCs in both *Pcna^R/R^* and *Rev1^-/-^*embryos compared to wildtype (**Fig. 4d**). In the soma however, we observed a comparable number of EdU^+^ somatic cells in mutant and wildtype embryos, suggesting that the reduced proliferation is restricted to the germ cell compartment (**Supplementary Fig. 8d**). To assess if reduced incorporation of EdU may be due to defects in cell cycle kinetics, we assessed the cell cycle properties of mutant PGCs. First, we stained cells for Ser-10 phosphorylation of histone H3 (pH3) that occurs during cell division facilitating chromatin compaction, necessary for mitosis (*64*). We found that a higher proportion of PGCs stained positive for pH3 in *Pcna^R/R^* and *Rev1^-/-^*genital ridges (**Fig. 4e**). We also looked at the localization of cyclin B1 which is localized in the cytoplasm during G2, before becoming phosphorylated during mitosis, driving its re-localization to the nucleus (*65, 66*). At E12.5 around 20% of wildtype PGCs had nuclear cyclin B1 and were therefore in G2/M-phase (**Fig. 4f**). This was significantly higher in the absence of TLS with an approximately 2-fold increase in G2/M-phase PGCs in *Pcna^R/R^* gonads (**Fig. 4f**). These data reveal that *Pcna^R/R^*and *Rev1^-/-^* PGCs accumulate markers of DNA damage, reduced proliferation, abnormal cell cycle kinetics and increased cell death. The combination of these cellular defects likely explains the profound reduction in the number of PGCs in the absence of TLS factors.

### *Pcna^R/R^* and *Rev1^-/-^* PGCs are developmentally blocked

Despite their scarcity, there are a few remaining PGCs in *Pcna^R/R^* and *Rev1^-/-^* embryos at E12.5 (medians *Pcna^R/R^*= 7 and *Rev1^-/-^* = 15). However, adult mice are completely sterile suggesting that the remaining PGCs are not able to generate functional gametes. Successful PGC development requires the activation of the germ cell transcriptional program coupled to global epigenetic changes (*67–69*). We therefore set out to test if *Pcna^R/R^* and *Rev1^-/-^* PGCs underwent these critical transcriptional and epigenetic processes. Initially, we performed gene expression analysis on E12.5 PGCs and found that both *Pcna^R/R^* and *Rev1^-/-^* PGCs expressed the early markers of PGC development *Nanos3* and *Prdm1* at comparable levels to wildtype (**Fig. 5a**). Conversely, expression of the later-stage marker *Mvh* was dramatically reduced in both mutants (**Fig. 5a**). We extended the gene expression analysis to additional germ cell specific genes in *Pcna^R/R^* E12.5 PGCs. Consistent with the previous data, genes normally expressed in the later stages of PGC development (*Dazl*, *Mili*, *Mael, Sycp3, Mov10l1, Hormad1* and *Brdt*) had reduced expression compared to wildtype which was not true for early-stage genes (*Stella* and *Fragilis*) (**Fig. 5b**). Therefore, E12.5 TLS-deficient PGCs transcriptionally resemble earlier stages of development.

**Fig. 5.**
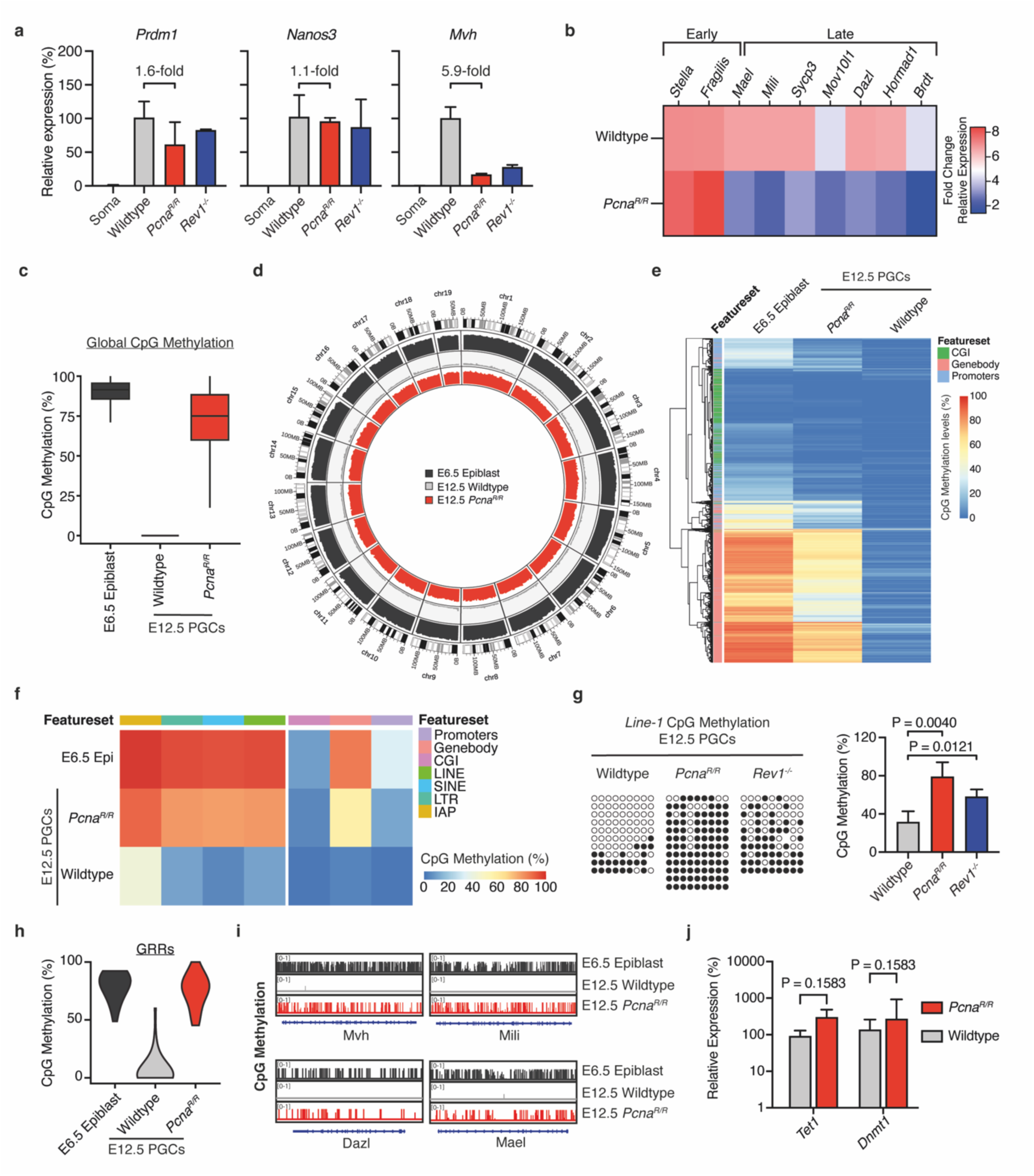
REV1 and PCNA K164 preserve the PGC developmental programme. **(a)** RT-qPCR expression analysis of early (*Prdm1* and *Nanos3*) and late (*Mvh*) germ cell markers in somatic cells and PGCs from wildtype, *Pcna^R/R^* and *Rev1^-/-^* E12.5 embryos (n=3 independent embryos per genotype). **(b)** RT-qPCR expression analysis of early (*Stella* and *Fragilis*) and late (*Dazl*, *Mili*, *Mael, Sycp3, Mov10l1, Hormad1* and *Brdt*) germ cell markers in PGCs from wildtype and *Pcna^R/R^* E12.5 embryos. (For *Stella*, *Fragilis*, *Dazl*, *Mili*, *Mael, Sycp3* and *Mov10l1*, wildtype n=6 and *Pcna^R/R^* n=7. For *Hormad1* and *Brdt*, wildtype n=3 and *Pcna^R/R^* n=5.) **(c)** Box plot of global DNA CpG methylation levels in wildtype E6.5 epiblast cells and E12.5 wildtype PGCs and *Pcna^R/R^* PGCs. Graph represents the median and interquartile range of CpG methylation distribution of the genome segmented in 5Kbp genomic windows. **(d)** Circos-plot representation of DNA methylation levels along the genome for E6.5 epiblast cells from wildtype embryos, E12.5 wildtype PGCs and E12.5 *Pcna^R/R^* PGCs. CpG methylation was averaged in 5 Mbp genomic windows and the average DNA methylation is represented as a histogram track. **(e)** Methylation heatmap showing methylation levels of CpG islands (CGI), gene bodies and gene promoters for E6.5 epiblast cells from wildtype embryos, E12.5 wildtype PGCs and E12.5 *Pcna^R/R^* PGCs. **(f)** Unclustered methylation heatmap representing levels of CpG methylation across CGIs, gene bodies, gene promoters and repeat elements in wildtype E6.5 wildtype epiblast cells and E12.5 wildtype and *Pcna^R/R^* PGCs. **(g)** Genomic bisulfite sequencing reads and qunaitifcation of the CpG-rich region of the *Line-1* element from FACS-purified somatic cells from E12.5 wildtype, *Pcna^R/R^* and *Rev1^-/-^* embryos (filled = methylated CpG, open = unmethylated CpG). **(h)** Violin plots reflecting the DNA methylation levels of GRR gene bodies in E6.5 epiblast cells from wildtype embryos, E12.5 wildtype PGCs and E12.5 *Pcna^R/R^* PGCs. The percentage methylation is calculated over each gene body with each point representing an individual gene. **(i)** Integrated genome viewer (IGV) visualisation of CpG methylation across selected GRR genes in E6.5 epiblast cells from wildtype embryos, E12.5 wildtype PGCs and E12.5 *Pcna^R/R^*PGCs. The plots represent the distribution of CpG methylation across genes segmented in 0.1 Kbp genomic windows. **(j)** RT-qPCR expression analysis of *Tet1* and *Dnmt1* in PGCs from wildtype and *Pcna^R/R^* E12.5 embryos. (For *Tet1*, n=5 for both genotypes. *Dnmt1*, wildtype n=6 and *Pcna^R/R^* n=7.)

A mechanism for the activation of germ cell specific gene expression in PGCs is through the DNA demethylation of their promoters (*14, 67–69*). This is linked to the process of epigenetic reprogramming in PGCs, specifically DNA demethylation, which occurs across the whole genome and is unique to the PGC compartment in the embryo. Alongside gene expression regulation, DNA demethylation is critical for imprint erasure and X-chromosome reactivation, processes needed for germ cell function. Our gene expression data prompted us to assess DNA methylation of PGCs. We performed whole genome bisulfite sequencing (WGBS) on wildtype and *Pcna^R/R^* FACS-purified E12.5 PGCs and compared these to wildtype E6.5 epiblast cells, which are the origin of PGCs (**Fig. 5c**). Compared to the epiblast cells the genome of wildtype E12.5 PGCs had extremely low levels of DNA CpG methylation, reflecting demethylation during PGC development. In contrast, in *Pcna^R/R^* PGCs we found a near complete retention of DNA CpG methylation (**Fig. 5c**). We mapped the methylation levels across the genome and found the retention in *Pcna^R/R^* PGCs was genome-wide (**Fig. 5d**).

DNA methylation does not proceed uniformly, with different genomic features reaching the lowest level of methylation at different times (*7, 14, 68, 69*). We therefore asked if only a subset of genomic features retained DNA methylation in TLS mutants or if the defect was truly global. Analysis of gene bodies and promoters showed that these were hypermethylated in *Pcna^R/R^* PGCs compared to wildtype, suggesting a failure in the early, global phase of demethylation (**Fig. 5e-f**). We also found that repeat elements (LINE-1, SINE elements, and endogenous retroviruses) retained methylation in *Pcna^R/R^* PGCs (**Fig. 5f & Supplementary Fig. 9a**). We validated this result by locus-specific, targeted bisulfite DNA sequencing of LINE-1 as it makes up ∼20% of the genome and again found increased methylation in both *Pcna^R/R^* and *Rev1* knockout PGCs (**Fig. 5g & Supplementary Fig. 10a-d**). We went on to analyze features which undergo DNA demethylation in the later wave and for which demethylation is both characteristic of germ cell development but also required for function. We found that imprinted differentially methylated regions (DMRs) were hypermethylated in E12.5 *Pcna^R/R^* PGCs, including both maternally and paternally methylated DMRs (**Supplementary Fig. 9b**). A failure to erase these features is a prerequisite for establishing the sex-specific methylation patterns that control genomic imprinting. Thus, this analysis reveals that the later wave of DNA demethylation also fails in the absence of TLS.

GCI promoters of genes associated with germ cell specific processes have been shown to retain DNA methylation until E11.5 and be demethylated in the later wave of PGC DNA demethylation (*14*). These genes are involved in meiosis and gamete generation and are only expressed in germ cells but silenced and methylated in somatic cells. We found that these promoters were hypermethylated in mutant PGCs (**Supplementary Fig. 9c**). We next analyzed CpG methylation of genes required for germ cell production whose transcriptional activation is concurrent with promoter demethylation during the later stages of PGC development (germline responsive genes - GRRs) (*69*). In agreement, CpG methylation analysis of a panel of 45 GRRs showed that these were largely demethylated in E12.5 wildtype PGCs (**Fig. 5h**). In *Pcna^R/R^* E12.5 PGCs however, these were hypermethylated. We confirmed this by performing locus-specific BS-Seq of the promoters of two GRR genes, *Mili* and *Dazl* (**Supplementary Fig. 10e-f**). It is striking that we observed both reduced expression of the GRR genes *Mvh*, *Dazl*, *Mili*, *Mael, Sycp3, Mov10l1, Hormad1* and *Brdt* and also hypermethylation of these loci in *Pcna^R/R^* PGCs (**Fig. 5b, 5I & Supplementary Fig. 9d**). This retention of DNA methylation in TLS mutants may explain the reduced activation of GRRs.

We compared the levels of DNA CpG methylation in mutant and wildtype E12.5 PGCs to the E6.5 epiblast. The retention of methylation across a panel of genomic features revealed that E12.5 *Pcna^R/R^* PGCs were more akin to the epiblast than to wildtype PGCs (**Fig. 5e-g**). Together, these data show that loss of TLS prevents global DNA demethylation in PGCs with retention occurring at early and late demethylated regions. Furthermore, E12.5 mutant PGCs appear trapped at an earlier stage of development with a methylome similar to E6.5 epiblast cells.

Next, we focused on understanding the basis for methylation retention in the absence of TLS and its effect on expression of germline genes. First, we studied the methylation machinery that normally ensures methylation patterns are inherited during cell division. Following DNA replication, the newly synthesized DNA strand is unmethylated, however DNMT1 is recruited to hemimethylated CpG dyads and transfers a methyl group to the unmethylated cytosine. We measured the expression of DNMT1 and found no difference between wildtype and *Pcna^R/R^* E12.5 PGCs suggesting that overexpression of the maintenance methyltransferase was not responsible for DNA hypermethylation in the absence of TLS (**Fig. 5j**). The enzyme TET1 has been shown to play important DNA demethylation dependent and independent roles in regulating the expression of genes critical for gamete formation. Notably, TET1 has previously been shown necessary for the transcriptional activation and maintenance of low levels of methylation at GRRs (*69*). We quantified the expression of *Tet1* in wildtype and *Pcna^R/R^* E12.5 PGCs and found no difference indicating that failure to activate the GRR genes is not mediated through *Tet1* expression (**Fig. 5j**).

Together, these data reveal that E12.5 TLS deficient PGCs resemble earlier stages of germ cell development transcriptionally and in their methylome. As DNA methylation regulates gene expression it is plausible that the retention observed in TLS mutants may contribute to the observed reduction in expression of germ cell genes. As well as failure of GRR demethylation, mutant PGCs also fail to erase imprinted DMR methylation, processes essential for normal germ cell function.

## Discussion

Declining birth rates and increasing infertility are drawing attention to environmental and genetic causes of human infertility. GWAS studies have identified the DDR as an important regulator of this, however, mechanistic studies have for the most part been lacking. One process implicated through these studies is TLS and the results of our current work reveal a crucial role during germline development. Our findings demonstrate that REV1 is required for both human PGCLC and mouse PGC development, indicating a conserved embryonic cause of infertility.

Alongside REV1, we show that REV7 and the modification site of PCNA, K164, are essential for PGC development. Though all three of these factors can act in multiple different processes, our results assert that it is their common role in TLS that is required (*29*). REV1 has both catalytic and non-catalytic functions and we have found that the C-terminal protein interaction domain, critical for TLS, is required in PGCs. Like REV1, the monoubiquitination of PCNA at K164 serves as a TLS scaffold. Importantly, the extender polymerase in TLS, Polζ, is recruited to TLS sites through direct interactions between its REV7 subunit and REV1’s C-terminal domain (*29, 35*). The finding that both the REV1 C-terminus and REV7 are essential in PGCs supports a role for both in TLS. Our results did not find a requirement for an inserter TLS polymerase. However, whilst Polζ has a specialized function in catalyzing DNA synthesis after lesion bypass, there is well-characterized redundancy between the inserter polymerases and hence this is not surprising (*18, 19*). Building on this, we found that the three core TLS factors are required during embryonic germ cell development and have identical timing and magnitude of PGC defects. In addition, the remaining PGCs in *Rev1^-/-^* and *Pcna^R/R^* embryos are phenotypically indistinguishable. This striking phenotypic overlap suggests that the cause of defect may be the same in all three mutants with TLS being the only function common to all three factors. Together, these data argue that TLS plays an essential role in fertility by preserving the development of PGCs.

The phenotypic features of the few remaining PGCs in mutant embryos explains the basis of germ cell loss and sterility. We found that mutant PGCs have a higher burden of DNA damage, apoptosis and cell cycle abnormalities as well as a failure in DNA demethylation and activation of the germ cell transcriptional program. The numerical PGC defect is at least partly explained by a combination of cell cycle arrest and increased apoptosis. However, defects in DNA demethylation and germline gene expression are also likely to significantly contribute to infertility, as defects in either of these processes result in catastrophic failure of PGC development and infertility (*4, 69*). Our results reveal that TLS is essential for the developmental processes that are needed for the production of mature germ cells.

The increased DNA damage, cell cycle abnormalities and increased apoptosis, whilst not previously shown in a physiological context, are consistent with known consequences of TLS deficiency or genotoxin exposure in tissue culture systems. However, the failure to undergo DNA demethylation or transcribe germ cell factors is entirely unexpected. Other DNA repair pathways such as Fanconi Anaemia DNA crosslink repair, have previously been shown to be necessary for PGC development (*23, 25, 70–72*). However, loss of these factors does not alter the expression of germline responsive genes or block DNA demethylation (*23*). This shows that whilst multiple DNA repair pathways are required for normal PGC development, they have distinct roles with different phenotypic outcomes.

Whilst wildtype PGCs have very low levels of DNA methylation by E12.5, we found that TLS-deficient PGCs have a global retention of DNA methylation. Indeed, we found that the methylation of TLS-deficient PGCs at E12.5 was similar to that seen in E6.5 epiblast cells. These data argue that the PGC precursors in the epiblast undergo *de novo* methylation before later undergoing demethylation during PGC development. The observed cell cycle defects could explain why demethylation is prevented in the absence of TLS. Cell cycle arrested PGCs may be unable to respond to time-dependent cell extrinsic cues, such as signaling molecules that instruct PGCs to proceed in their development. Alternatively, the cell cycle perturbations may block a cell-intrinsic process, preventing activation of the germ cell program and epigenetic reprogramming. Interestingly, the first stage of global DNA demethylation in PGCs relies upon replication to dilute methylation marks (*14, 69*). Therefore, the cell cycle arrest in TLS mutant PGCs may result in the genome-wide retention of methylation marks that we observe. The normal expression of TET1 in mutant PGCs argues against dysregulation of either its enzymatic or non-enzymatic functions as a contributing factor to the phenotype observed.

Whilst early stage PGC genes are induced by extrinsic signals and maintained by PGC-specific and pluripotency-associated transcription factors, DNA demethylation is important for the transcription of genes expressed in the later stages of PGC development. Therefore, the failure to transcribe GRR genes and retention of DNA methylation may not be independent of each other. Indeed, we find reduced mRNA expression and DNA hypermethylation of each GRR gene that was analyzed. It is therefore likely that the block in cell cycle progression in the absence of TLS prevents DNA demethylation, transcription of germ cell factors and therefore blocks PGC development.

The specificity of the phenotype which only affects germ cell homeostasis is intriguing. *In vivo* assessment of rapidly dividing tissues in *Pcna^R/R^* mice revealed no defects in somatic tissue homeostasis. Despite a reduction in blood stem cell frequency that we show begins *in utero*, TLS-deficient mice sustain peripheral blood homeostasis throughout life. Moreover, in the absence of TLS, mice have normal longevity and no overt phenotypes in other somatic tissues. In contrast, there is a complete failure to generate functional gametes. This striking dichotomy raises the question of why TLS is only required to maintain homeostasis in the germ cell compartment. We have demonstrated that TLS is required for processes that occur only in the germ cell lineage: engagement of germline genes and genome-wide DNA demethylation. In addition, we find that the canonical phenotypes associated with TLS deficiency and the DNA damage response are also restricted to the germ cell compartment: unresolved DNA damage, apoptosis and cell cycle perturbation (**Fig. 4 and Supplementary Fig. 8**). This raises the possibilities that i) the DNA damage checkpoints that activate apoptosis or cell cycle arrest are different in PGCs than in somatic cells, ii) there are replication impediments or other features unique to early PGCs that confer a dependency on TLS activity or iii) that PGCs are more reliant upon TLS to resolve replication impediments than somatic cell types. Finally, it is tempting to speculate that the dependency of PGC development on TLS may be connected to the early epigenetic reprogramming events required for subsequent germline development.

**Table.1.** *PcnaK164R* mice are born at the expected ratios. *Pcna^R/R^* embryos are observed at the expected Mendelian ratios throughout development (E11.5 and E12.5) and at 21 days postpartum (P21).

|  | *Pcna*<sup>+/+</sup> | *Pcna*<sup>+/*R*</sup> | *Pcna*<sup>*R/R*</sup> |  |
| --- | --- | --- | --- | --- |
| Exp. | **25%** | **50%** | **25%** |  |
| Obs. | 10.0%<br>n = 2/20 | 65%<br>n = 13/20 | 25%<br>n = 5/20 | E11.5 |
| Obs. | 31.8%<br>n = 27/85 | 45.9%<br>n = 39/85 | 22.4%<br>n = 19/85 | E12.5 |
| Obs. | 26.0%<br>n = 51/196 | 49.5%<br>n = 97/196 | 24.5%<br>n = 48/196 | P21 |

## Acknowledgments

We thank A. Surani for the gift of Stella-GFP mice. The hiPSC cell line (BTAG 585b1-868) was a kind gift from Prof. Mitinori Saitou (Institute for the Advanced Study of Human Biology, Kyoto, Japan). We thank the Human Research Tissue Bank (National Institute for Health Research Cambridge Biomedical Research Centre) for processing the histology. We would like to thank Keith Burling and Peter Barker of the NIHR Cambridge Biomedical Research Centre Core Biochemical Assay Laboratory for carrying out the biochemical assays for LH, FSH, albumin, AST, creatinine and urea.

## Funding

P.S., R.J.H. and G.P.C. are supported by the Medical Research Council as part of UK Research and Innovation (file reference no. MC_UP_1201/18)..

## Author contributions

G.P.C. conceived the study, designed experiments and wrote the manuscript. P.S. and R.J.H. designed and performed all experiments and wrote the manuscript.

C.D. generated mutant hiPSCs and performed hPGCLC assays. S.C. performed bioinformatic analysis of WGBS data. A.A. performed library prep for WGBS. M.J.A. analyzed pathological features of histology samples. H.L. and W.R. helped design experiments and write the manuscript.

## Competing interests

S.C., A.A., and W.R. are employees of Altos Labs.

## Data and materials availability

All data are available in the main text or the supplementary materials. All materials are available on request from the corresponding author.

## Materials and Methods

### Human iPSC culture

The hiPSC cell line (BTAG 585b1-868) was a kind gift from Prof. Mitinori Saitou (Institute for the Advanced Study of Human Biology, Kyoto, Japan). Cells were maintained in Stemfit Basic04 (Amsbio, SFB-504-CT) on Geltrex (Life Technologies, A1413302) coated dishes. Cells were passaged as single cells using Accutase (Millipore, SF006) and 10 μM of rock inhibitor (Y-27632, STEMCELL Technologies, 72304) was added for the first 24 hours after seeding. Cells were maintained in hypoxic conditions (5% CO_2_, 5% O_2_) and without antibiotics.

### Induction of iMeLCs and hPGCLCs

Induction of iMeLCs and hPGCLCs was performed as previously published by Sasaki et al (*26*). For iMeLC induction, hiPSCs were dissociated as single cells using a 1:1 mixture of TrypLE (Life Technologies, 12604013) and EDTA/PBS. 1.5 × 10^5^ cells were seeded on a fibronectin coated well of a 12 well plate in GK15 medium (GMEM [Life Technologies] with 15% KSR, 0.1 mM NEAA, 2 mM L-glutamine, 1 mM sodium pyruvate, and 0.1 mM 2-mercaptoethanol) supplemented with: 50 ng/μl of ActivinA (R&D, 338-AC-010), 3 μM of CHIR99021 (Cambridge Bioscience, SM13-5) and 10 μM of Y-27632. For hPGCLCs, 3.0 × 10^3^ iMeLCs were plated into ulta-low attachment U-bottom 96-well plate (PHCBI, MS-9096UZ) in GK15 supplemented with: 10 ng/ml LIF (Millipore, LIF1005), 200 ng/ml of BMP4 (314-BP-010, 100 ng/ml of SCF (R&D Systems, 255-SC-050), 50 ng/ml of EGF (R&D Systems, 236-EG-200), and 10 μM of Y-27632.

### Generation of *Rev1^-/-^* BTAG hiPSC line

Two gRNAs were designed to target the exon 3 of *REV1* (**Supplementary Table 2**) and subcloned into the PX458 plasmid (Addgene, 48138). 5.0-8.0 × 10^5^ BTAG cells were electroporated with 3 μg of each gRNA using Amaxa 4D nucleofector (Lonza, V4XP-3012) and the CA-137 program. 48 hours after the transfection, GFP positive cells were FACS sorted and plated in 96 well plate with one cell per well in Stemfit Basic04, 10μM of Y-27632, penicillin and streptomycin (Life Technologies, 15070063). After 10-14 days single cell-derived colonies were picked and expanded. Clones were screened by PCR on extracted genomic DNA (NEB, T3010L) using the primer pair listed in Supplementary Table 2.

### Mitomycin C (MMC) treatment of BTAG hiPSC cells

Wildtype and *Rev1^-/-^* cells were passaged as single cells then seeded at 1:10 density in Stemfit Basic04 and Y-27632. 24 hours after, medium was changed and Y-27632 was removed. The next day, 5 ng/ml MMC (Insight Biotechnology, sc-3514A) was added to the medium and 48-72 hours later wells were assessed for surviving hiPSC colonies.

### hPGCLC immunostaining

Day 4 hPGCLC aggregates were fixed in 4% PFA for 20 min at room temperature. After PBS washes and incubation in 30% sucrose, aggregates were embedded in OCT (VWR, 361603E) and sectioned at a thickness of 10 μm. Samples were permeabilised for 20 min in PBS/0.2% Triton X100 (PBST) and blocked for 1 hour in blocking solution (Santa Cruz, sc-516214). Samples were then incubated with the following primary antibodies diluted in blocking buffer at room temperature for 2 hours: anti-OCT4 (1:200, catalog no. ab181557; Abcam); anti-TFAP2C (1:100, catalog no. sc-12762; Santa Cruz); anti-SOX17 (1:200, catalog no. AF1924; R&D). After PBS washes, samples were incubated with the following secondary antibodies diluted 1:1000 in PBS: Donkey anti-rabbit IgG (Alexa Fluor 594, catalog no. A21207; Invitrogen); Donkey Anti-Goat IgG H&L (Alexa Fluor 488, catalog no. ab150129; Abcam); Donkey anti-Mouse IgG (H&L) (Alexa fluor 594, catalog no. R37115; Life Technologies). Sections were then washed with PBS and incubated for 10 min with DAPI. A coverslip was then placed on the slides in Vectashield vibrance (Vector Labs, H-1800-10). Sections were imaged using the SP5 inverted confocal microscope (Leica).

### Flow cytometry analysis of hPGCLC aggregates

Day 4 aggregates were collected and dissociated with 0.25% trypsin-EDTA (Life Technologies, 25200056) for 10 min at 37℃ under agitation. Cells were washed with PBS containing fetal bovine serum (FBS) and 0.1% bovine serum albumin (BSA) before being subjected to centrifugation. Dissociated cells were then resuspended in FACS buffer (PBS, 0.1% BSA), filtered by a cell strainer (BD Biosciences) and analysed or sorted by FACS (Aria III, BD Biosciences) based on the eGFP and tdTomato reporters. Analysis of hPGCLCs differentiation efficiency was performed on one wild type clone and three REV1 KO clones in at least three independent experiments. The cytometry software FlowJo™ version 10 was used for analysis.

### Mice

All animal experiments performed in this study were approved by the Medical Research Council’s Laboratory of Molecular Biology animal welfare and ethical review body and conform to the UK Home Office Animal (Scientific Procedures) Act 1986 (License no PP6752216). All mice were maintained under specific pathogen-free conditions in independently ventilated cages (GM500; Techniplast) on Lignocel FS-14 spruce bedding (IPS) and provided with environmental enrichment (fun tunnel, chew stick and Enviro-Dri nesting material (LBS)) at 19-23°C. Mice fed Dietex CRM pellets (SpecialDietService) ad libitum with light from 7:00 a.m. to 7:00 p.m. No mice used in this study were wild and no field-collected samples were used. All mice were maintained on an isogenic C57BL/6j background. Embryos were examined at various developmental stages from E8.5 to E13.5 as indicated in the text. Female mice used in timed-mating experiments were aged between 6-25 weeks. The investigators were blinded to the genotypes of all mice throughout the study and data were acquired by relying entirely on identification numbers. The *Stella-GFP* (*Tg(Dppa3/EGFP)6-25Masu*) allele (MGI ID: 5519126) was a kind gift from Azim Surani. B6;129P2-*Polk^tm1.1Rsky^/J* allele (MGI ID: 2445458) and the B6.Cg-*Polq^tm1Jcs^/J* allele (MGI ID: 2155399) have been described previously. *GOF18*-*GFP*(*Tg(Pou5f1-EGFP)2Mnn*) (MGI ID: 3057158) JAX (stock ID: 004654) mice were purchased from The Jackson Laboratory.

### Generation of *Pcna^K164R^*, *Rev1^AA^* and *Rev1^CT^* mutant mice

Mice carrying the *Pcna^K164R^* and *Rev1^CT^* alleles were generated by Alt-R CRISPR-Cas9 (IDT) mediated genome editing in zygotes on a C57BL/6J background. tracrRNA and crRNA were diluted to a final concentration of 1μg/ul in injection buffer (10 mM Tris HCl pH 7.5, 0.1 mM EDTA). 5 μg of crRNA and 10 μg tracrRNA were mixed and annealed by heating to 95°C for 5 min and then ramped to 25°C at a rate of 5°C per min. Alt-R SpCas9-3NLS was diluted to a concentration of 200 ng/μl in injection buffer. The RNP was assembled by diluting Alt-R SpCas9-3NLS and the annealed crRNA:tracrRNA at a final concentration of 20 ng/μl and incubated at room temperature for 15 min. The ssODN was then added to the RNP complex at a final concentration of 20 ng/μl and injected into zygotes. The zygotes were surgically transplanted into pseudopregnant CD1 females. Progeny were screened for correct gene targeting and the targeted allele sequenced. *Rev1^AA^* mice were generated by gene targeting in mouse embryonic stem cells. Stella-GFP Bac9 ESCs were transfected with an ssODN and px451 plasmid containing a guide targeting exon 10 of *Rev1*. Clones were screened by PCR followed by AciI restriction digest. Two positive ESC clones were injected into C57BL6/J:TYR blastocysts. Mice with high levels of chimerism were back crossed and the progeny screened by PCR.

### Isolation of *Pcna^K164R^*, *Rev1^AA^* and *Rev1^CT^* mouse embryonic fibroblasts

Timed matings were performed between heterozygous mice carrying either the *Pcna^K164R^*, *Rev1^AA^* or *Rev1^CT^*mutations. Pregnant females were culled at E12.5 to harvest embryos. Embryos were incubated in pre-warmed trypsin solution (2.5 μg.mL^-1^ trypsin (Gibco), 25 mM Tris, 120 mM NaCl, 25 mM KCl, 25 mM KH_2_PO_4_, 25 mM glucose, 25 mM EDTA, pH 7.6) for 10 min and disaggregated by gentle pipetting. Primary mouse embryonic fibroblast (MEF) cultures were established following standard methods and immortalized using the SV40 large T antigen as described previously. Briefly, Platinum-E retroviral packaging cells (Cell Biolabs) were transfected with pBABE-SV40-Puro and the culture media containing the virus was harvested 48 hours later and passed through a 0.22 μm filter. The filtered retrovirus was mixed 1:1 with complete MEF media supplemented with 1 μg.mL^-1^ hexadimethrine bromide (Polybrene, Millipore). The infective medium was subsequently added to primary MEF cultures and transformed clones were selected for 14 days using 1 μg.mL^-1^ puromycin.

### Measuring sensitivity of MEFs to DNA damaging agents

Sensitivity to DNA-damaging agents was determined by seeding 1,000 transformed MEFs per well of a 96-well flat-bottom plate and exposing to ultraviolet (*57*) irradiation or mitomycin C (MMC). After 7 days of culture the MTS cell viability reagent (CellTiter 96^®^ Aqueous One Solution Cell Proliferation Assay, Promega) was added and plates incubated for 4 hours at 37°C; absorbance was then measured at 492 nm.

### Histological analysis

Tissues were fixed in 10% neutral-buffered formalin for 24-36 hours then transferred to 70% ethanol. Fixed samples were dehydrated and embedded in in paraffin and 4 μm sections cut. Sections were deparaffinised, re-hydrated and stained with haematoxylin and eosin (H&E) following standard methods. Images were captured with an Eclipse Ti2-E (Nikon) microscope and tissue architecture was scored blindly.

### Immunohistochemistry

Formalin-fixed, paraffin-embedded samples were sectioned at 4 μm, deparaffinised and rehydrated following standard methods. Slides were boiled in antigen retrieval buffer (10 mM sodium citrate, pH 6) for 10 min and allowed to cool to room temperature before being washed three times in water for 5 min and then once in TBS, 0.1% w/v Tween-20 for 5 min. A hydrophobic ring was drawn around the tissue sections and samples incubated with blocking buffer (TBS, 0.1% w/v Tween-20, 5% v/v goat serum) for 1 hour at room temperature. Samples were incubated with the following primary antibodies diluted in blocking buffer at 4°C overnight: anti-PLZF (1:200, sc-28319) and anti-Ki67 (1:200, ab16667). Slides were washed three times with TBS, 0.1% w/v Tween-20 for 5 min and incubated with the following secondary antibodies for 1 hour at room temperature: swine anti-rabbit (1:200, catalog no. P0339, Dako) or anti-mouse horseradish peroxidase (HRP)-conjugated immunoglobulins (1:200, catalog no. P0339, Dako). For HRP-based immunohistochemistry, slides were incubated for 3-10 min with SignalStain diaminobenzidine substrate kit (catalogue no. 8059P; Cell Signalling Technology) and then washed once in water for 5 min. Slides were dehydrated in an ethanol gradient following standard methods and finally in xylene before being mounted with DPX neutral mounting medium (catalog no. 317616, Sigma-Aldrich) and coverslips place on top of the slides. Images were captured with an Eclipse Ti2-E (Nikon) microscope. The frequency of PLZF^+^ or Ki67^+^ cells were scored blindly.

### Assessing fertility of mice

Mice were paired with wildtype C57BL/6J mice of the opposite sex. Female mice were monitored daily for the presence of copulation plugs and the number of offspring born over three successive months was recorded (only data from breeding pairs where at least 3 copulation plugs were observed was included in the analysis). Investigators performing the copulation plug checks were blinded to the genotypes of the mice.

### Timed matings for embryo isolation

Timed matings were performed overnight and female mice were assessed for the presence of copulation plugs the following day and separated from males. Halfway through the light cycle on the day a copulation plug was observed was designated E0.5. Pregnant mice were culled at noon of the appropriate day during gestation (E8.5–13.5) and the embryos harvested. Samples were processed immediately for further analysis with a small tissue biopsy taken for genotyping.

### Immunofluorescence

Timed matings were performed as described above and E12.5 embryos harvested. The fetal gonads were dissected out and fixed in PBS, 4% w/v paraformaldehyde for 30 min at 4°C. Fixed samples were washed three times in PBS, 1% w/v Triton X-100 for 5 min at room temperature for permeabilization then pressed onto glass slides. A large hydrophobic ring was drawn around each sample and then incubated in blocking buffer (PBS, 1% w/v Triton X-100, 1% w/v BSA) for 1 hour at room temperature. Samples were then incubated with the following primary antibodies at 4°C overnight: anti-GFP (1:500, catalog no. GF090Rl; Nacalai); anti-GFP (1:2,000, catalog no. ab13970); anti-phospho-Histone H2A.X (Ser139) (1:1,000, catalog no. 05-636; Millipore); anti-cleaved caspase 3 (1:400, catalog no. 9661; Cell Signaling Technology); anti-MVH (ab27591); anti-RPA (1:100, catalog no, 2208; Cell Signalling Technology); anti-cyclin B1 (1:100, catalog no. 4138; Cell Signalling Technology); anti-phospho-histone 3 (1:200, catalog no, 9701; Cell Signalling Technology). Samples were then washed three times in PBS, 1% w/v Triton X-100 for 5 min and then incubated for 1 hour at room temperature with the following secondary antibodies: goat anti-rat Alexa Fluor 488 (1:1000, catalog no. A11006, Thermo Fisher Scientific), goat anti-chicken Alexa Fluor 488 (1:1000, catalog no. A11039, Thermo Fisher Scientific), goat anti-mouse Alexa Fluor 594 (1:1000, catalog no. A11032, Thermo Fisher Scientific), goat anti-rabbit Alexa Fluor 594 (1:1000, catalog no. A21429; Thermo Fisher Scientific), goat anti-rat Alexa Fluor 594 (1:1000, catalog no. A21209; Thermo Fisher Scientific). Slides were then washed three times in PBS, 1% w/v Triton X-100 for 5 min and stained with 0.5 μg.mL^-1^ DAPI diluted in PBS for 10 min. Slides were washed once then mounted with ProLong Gold Antifade mounting media (catalog no. P36934; Thermo Fisher Scientific). Coverslips were placed on top of the slides and the slides were allowed to cure for 48 hours. Images were captured with an LSM780 confocal microscope (Zeiss) and specimens were scored blindly.

### Preparation of PGCs and their quantification and isolation by flow cytometry

For PGC quantification, the entire embryo (E8.5-10.5) or the developing urogenital ridges (E11.5-13.5) were isolated from embryos and placed into 500 μL or 150 μL pre-warmed trypsin solution (2.5 μg.mL^-1^ trypsin (Gibco), 25 mM Tris, 120 mM NaCl, 25 mM KCl, 25 mM KH_2_PO_4_, 25 mM glucose, 25 mM EDTA, pH 7.6) respectively, and incubated at 37°C for 10 min. Subsequently, 1 μL of Benzonase endonuclease (Millipore) was added and the sample gently disaggregated by pipetting and incubated for 5 min at 37°C. The trypsin was inactivated by adding 1 mL of PBS/5% v/v fetal bovine serum and centrifuged at 3,300 rpm for 10 min. The sample was resuspended in 100 μL of anti-SSEA1 conjugated to Alexa Fluor 647 (catalog no. MC-480; Biolegend) diluted 1:100 in PBS/2.5% v/v fetal bovine serum and incubated for 10 min at room temperature. Samples were diluted by adding 300 μL of PBS/2.5% v/v fetal bovine serum and passed through a 70 μm filter. For quantification, 300 μL of the samples were immediately run on an ECLIPSE analyzer (Sony Biotechnology) and the data analyzed using FlowJo v10.1r5. For sorting of cells, samples were immediately run on a Synergy cell sorter (Sony Biotechnology), the cells sorted into 10 μL of PBS, centrifuged at 6,000 r.p.m. for 5 min and stored at -80°C until further analysis.

### Gene expression analysis

The expression of *Pcna*, *Rev1*, *Polk* and *Polq* in PGCs was assessed by droplet digital PCR (ddPCR). PGCs (GOF18-GFP^+^SSEA1^+^) and somatic cells (GOF18-GFP^-^SSEA1^-^) were isolated by FACS as described above. Total RNA was extracted using the PicoPure RNA Isolation Kit (Thermo Fisher Scientific) and first-strand complementary DNA was synthesized using the SuperScript IV Reverse Transcriptase (Thermo Fisher Scientific) according to the manufacturer’s instructions. ddPCR was performed using ddPCR Supermix for Probes (BioRad) on a QX ONE Droplet Digital PCR (BioRad) following the manufacturer’s instructions. Taqman probes were purchased from ThermoFisher; Pcna, Rev1, Polk, Polq labelled with FAM and Gapdh with VIC. The expression was normalised to *Gapdh* and made relative to the somatic cells. The expression of *Rev1* in MEFs was assessed by purifying RNA with the RNeasy kit (Qiagen) according to the manufacturer’s instructions and first-strand complementary DNA was synthesized using the SuperScript IV Reverse Transcriptase (Thermo Fisher Scientific) according to the manufacturer’s instructions. PCR amplification was performed using the TaqMan Fast Advanced Master Mix (Thermo Fisher Scientific). PCR amplification was performed on a ViiA 7 cycler for 95 °C for 15 s and 60 °C for 1 min. Mean threshold cycles were determined from three technical repeats using the comparative CT methodology. All expression levels were normalized to *Gapdh* (mm99999915_g1). To perform gene expression analysis on *Rev1^-/-^* and *Pcna^R/R^* PGCs, 50 cells were sorted into 10 μL of Single cell lysis buffer supplemented with DNAseI (Single Cell-to-CT Kit ThermoFisher, 4458237) and stored at -80°C. The lysis and reverse transcription was performed using the single cell-to-CT kit (ThermoFisher, 4458237) according to the manufacturer’s description. Pre-amplification was performed with Taqman Probes (Thermofisher) targeting *Ddx4*, *Nanos3*, *Prdm1* and *Gapdh*. PCR amplification was performed using the TaqMan Fast Advanced Master Mix (Thermo Fisher Scientific). PCR amplification was performed on a ViiA 7 cycler for 95 °C for 15 s and 60 °C for 1 min. Mean threshold cycles were determined from three technical repeats using the comparative C_T_ methodology. All expression levels were normalized to *Gapdh*. For expression analysis of *Stella*, *Fragilis*, *Dazl*, *Mael*, *Mili, Sycp3, Mov10l1, Dazl, Hormad1* and *Brdt* E12.5 wildtype and *Pcna^R/R^* PGCs were FACS purified as described above. Subsequently, RNA was prepared using the NEBNext^®^ Single Cell/Low Input RNA Library Prep Kit for Illumina^®^ kit and NEBNext^®^ Poly(a) mRNA Magnetic Isolation module kit following the manufacturer’s instructions. Subsequently, all libraries were ligated to NEBNext^®^ Multiplex Oligos for Illumina^®^ (Dual Index Primer Set I) and quality control was performed using a 2100 Bioanalyser High Sensitivity DNA Kit 5067-4626 (Agilent) and libraries quantified using a Qubit^TM^ Fluorometer following the manufacturer’s instructions. Finally, RNA sequencing libraries were pooled to a final concentration of 8.5 nM. RNA expression was performed using Brilliant II SYBR Green QPCR Master Mix (Agilent) using a ViiA 7 Real-Time PCR system (Thermo Fisher Scientific) at 95°C for 10 min and 40 cycles of 95°C for 15 s and 60°C for 1 min. Mean threshold were determined from three technical repeats per sample and oligonucleotide pair using standard comparative C_T_ methods. All expression levels were normalized to *Gapdh*. For the testis specific genes *Mov10l1* and *Brdt*, gene expression values were only calculated for male samples.

### Bisulfite Sequencing

PGCs (GOF18-GFP^+^SSEA1^+^) and somatic cells (GOF18-GFP^-^SSEA1^-^) were isolated from E12.5 embryos carrying the GOF18-GFP reporter by FACS as described above. Genomic DNA extraction and sodium bisulfite conversion was performed using the EZ DNA Methylation-Direct Kit (Zymo Research Cat. No. D5020) following the manufacturer’s instructions. Nested oligonucleotide pairs described previously were used to amplify *Line-1* (*6*), *Dazl* (*68*) and *Mili* (*68*) sequences using the ZymoTaq polymerase (Zymo Research). PCR products were separated out on an agarose gel and gel-purified using the QIAquick Gel Extraction Kit (Qiagen) following the manufacturer’s instructions. Gel extracted products were ligated into the pGEM-T Easy Vector System I (Promega). Ligation products were transformed into *E. coli* and single clones picked following blue-white selection and sent for sequencing using M13R primer. Sequencing reads were analyzed using Quantification for Methylation Analysis (QUMA) software with standard quality control settings.

### Whole Genome Bisulfite Sequencing

PGCs (GOF18-GFP^+^SSEA1^+^) and somatic cells (GOF18-GFP^-^SSEA1^-^) were isolated from E12.5 embryos carrying the GOF18-GFP reporter by FACS as described above. WGBS-sequencing libraries were prepared according to the bulk version of the published scBS-seq protocol (https://www.nature.com/articles/nprot.2016.187). Briefly, cells were lysed in RLT plus buffer (Qiagen) then bisulfite converted using the Zymo EZ-Direct kit. Converted DNA was prepared for Illumina sequencing via two rounds of random primed synthesis using oligos containing Illumina adapter sequences followed by PCR to amplify and incorporate indexes. Sequencing was performed on a NextSeq 500 instrument using 75bp paired end reads. Bismark v0.23.1 (https://doi.org/10.1093/bioinformatics/btr167) was used to align DNA reads to the bisulfite converted GRCm38 mouse genome then perform methylation calling. E6.5 Epiblast data was taken from our previous work (https://doi.org/10.1038/s41586-019-1825-8) selecting a random subset of 50 E6.5 Epiblast cells. For analysis of genomic features (e.g. promoters, gene bodies, 5Kbp tiles), CpG methylation rates were computed assuming a binomial model as previously (https://doi.org/10.1038/s41586-019-1825-8). To generate the Circos plot, DNA-methylation bigwig files were read into python using pyBigWig library and resampled to 5Mbp bins. The chromosome band data is obtained from the UCSC table “cytoBandIdeo” of the GRCm38/mm10 assembly. The Circos plot was generated using the pycircos library.

### EdU incorporation analysis *in vivo*

To assess the *in vivo* incorporation of 5-ethynyl-2’-deoxyuridine (EdU) into the DNA of PGCs (GOF18-GFP^+^) the Click-iT™ Plus EdU Cell Proliferation Kit for Imaging, Alexa Fluor™ 594 kit (Catalog number: C10639) was used. Pregnant mice were given a single dose of EdU (50 mg.kg^-1^) by intraperitoneal injection (IP) at 10 ml.kg^-1^. Females were subsequently culled 4 hours post-IP at E12.5 and the embryos harvested; the fetal gonads were dissected and placed into ice-cold PBS. Fetal gonads were fixed in PBS, 4% w/v paraformaldehyde for 30 min at 4°C. Fixed samples were washed once in PBS for 5 min and then three times in PBS, 1% w/v Triton X-100 for 15 min at room temperature. Samples were then pressed onto glass slides and a large hydrophobic ring drawn around the sample before being incubated in blocking buffer (PBS, 1% w/v Triton X-100, 1% w/v BSA) for 1 hour at room temperature. Samples were then incubated with anti-GFP (1:500, catalog no. GF090R; Nacalai) diluted in blocking buffer overnight at 4°C. Subsequently, samples were washed three times in PBS, 1% w/v Triton X-100 for 5 min at room temperature and fixed in PBS, 2% w/v paraformaldehyde for 20 min at room temperature. Slides were washed three times in PBS and incubated in 200 μL of Click-iT® Plus reaction cocktail made per manufacturer’s instructions. Samples were washed three times in PBS, 1% w/v Triton X-100 and incubated with goat anti-rat Alexa Fluor 488 (1:1000, catalog no. A11029, Thermo Fisher Scientific) secondary antibody. Slides were washed three times in PBS, 1% w/v Triton X-100 for 5 min and stained with DAPI (0.5 μg.mL^-1^, PBS) before being mounted with ProLong Gold Antifade Mountant (catalog no. P36934; Thermo Fisher Scientific). Coverslips were placed on top of the slides and allowed to cure for 48 hours before images were captured with an LSM780 confocal microscope (Zeiss) and the frequency of positive cells scored blindly.

### Blood counts

A 50 μL total blood sample was taken from saphenous veins or via cardiac puncture and transferred into a K3EDTA MiniCollect tubes (Greiner bio-one) and analysed on a VetABC analyzer, using standard settings for mice (Horiba).

### Serum hormone analysis

For the determination of serum luteinizing hormone, follicle stimulating hormone (FSH) and testosterone levels, blood samples were collected as described above and transferred into a Microvette collection tube (SARSTEDT) and centrifuged at 10,000 r.p.m. for 10 min at room temperature. The supernatant (serum) was transferred to a 1.5 mL eppendorf tube and stored at -80°C until further analysis. For serum LH and FSH concentrations were determined using the Milliplex Map Mouse Pituitary Magnetic Bead Panel (catalog no. MPTMAG-49K). Plates were loaded manually and washed using a Bio-Plex Pro wash station (BioRad) and data analysis performed using a Magpix Multiplexer reader (BioRad). Serum testosterone levels were determined using the Demeditec Diagnostics rat/mouse ELISA kit (catalog no. DEV9911). Samples were loaded manually and washes performed using a WellWash Versa platewasher (Thermo Scientific). Absorbance was measured at 450 nm using Perkin Elmer Multicalc software. Serum levels of urea, creatinine, aspartate aminotransferase and albumin were measured using a Siemens Dimension RxL analyser.

### Haematopoiesis Analysis

Flow cytometry was performed on bone marrow cells that were isolated from the femora and tibiae of mutant mice and appropriate controls by flushing cells and passing them through a 70-μm filter. The following antibodies were used to stain for HSCs: FITC-conjugated lineage cocktail with antibodies anti-CD4 (clone H129.19, BD Pharmingen), CD3e (clone 145-2C11, eBioscience), Ly-6G/Gr-1 (clone RB6-8C5, eBioscience), CD11b/Mac-1 (clone M1/70, BD Pharmingen), CD45R/B220 (clone RA3-6B2, BD Pharmingen), Fcε R1α (clone MAR-1, eBioscience), CD8a (clone 53-6.7, BD Pharmingen), CD11c (clone N418, eBioscience) and TER-119 (clone Ter119, BD Pharmingen), anti-c-Kit (PerCP-Cy5.5, clone 2B8, eBioscience), anti-Sca-1 (PE-Cy7, clone D7, eBioscience). When staining for SLAM markers the same lineage cocktail was used (FITC) with the addition of the following antibodies: anti-CD48 (FITC, clone HM48-1, BioLegend), anti-CD41 (FITC, clone MWReg30, BD Pharmigen), anti-CD150 (APC, clone TC15-12F12.2, BioLegend) and anti-c-Kit and Sca-1 as above. Maturation of B cells was assessed using anti-CD45R/B220 (PE, clone RA3-6B2, BD Pharmingen) and anti-IgM (APC, clone II/41, BD Pharmingen). The maturation of the erythroid lineage was analysed using antibodies anti-TER-119 (APC, clone Ter-119, BD Pharmingen) and anti-CD71 (PE, clone C2, BD Pharmingen). Granulocyte–macrophage maturation was assessed with antibodies anti-CD11b/Mac-1 (APC, clone M1/70, BD Pharmingen) and anti-Ly-6G/Gr-1 (PE, clone RB6-8C5, eBioscience). Thymic T-cell maturation was assessed using CD4 (FITC, clone H129.19, BD Pharmingen) and CD8a (PE, clone 53-6.7, BD Pharmingen) antibodies. The samples were incubated for 15 min at 4 °C in the dark with the exception of samples containing anti-CD34 (RAM34), which were incubated for 90 min. Samples were run on a LSRII flow cytometer (BD Pharmingen) and the data were analysed with FlowJo 9.3.1 (Tree Star).

### Statistical Analysis

The number of independent biological samples and technical repeats (*n*) are indicated in figure legends. Unless otherwise stated, data are shown as mean ± standard deviation (s.d.) and the nonparametric Mann-Whitney test was employed to determine statistical significance. Analysis was performed in GraphPad Prism version 8.

## Notes

### Competing Interest Statement

The authors have declared no competing interest.

## References

1. M. Saitou, M. Yamaji, Primordial germ cells in mice. Cold Spring Harb Perspect Biol 4, (2012).

2. P. Soriano, R. Jaenisch, Retroviruses as probes for mammalian development: allocation of cells to the somatic and germ cell lineages. Cell 46, 19–29 (1986).

3. H. Ueno, B. B. Turnbull, I. L. Weissman, Two-step oligoclonal development of male germ cells. Proc Natl Acad Sci U S A 106, 175–180 (2009).

4. M. Saitou, S. C. Barton, M. A. Surani, A molecular programme for the specification of germ cell fate in mice. Nature 418, 293–300 (2002).

5. Y. Yabuta, K. Kurimoto, Y. Ohinata, Y. Seki, M. Saitou, Gene expression dynamics during germline specification in mice identified by quantitative single-cell gene expression profiling. Biol Reprod 75, 705–716 (2006).

6. P. Hajkova et al., Epigenetic reprogramming in mouse primordial germ cells. Mech Dev 117, 15–23 (2002).

7. N. B. Ramakrishna, K. Murison, E. A. Miska, H. G. Leitch, Epigenetic Regulation during Primordial Germ Cell Development and Differentiation. Sex Dev 15, 411–431 (2021).

8. Y. Zeng, T. Chen, DNA Methylation Reprogramming during Mammalian Development. Genes (Basel) 10, (2019).

9. P. Hajkova et al., Chromatin dynamics during epigenetic reprogramming in the mouse germ line. Nature 452, 877–881 (2008).

10. Y. Seki et al., Cellular dynamics associated with the genome-wide epigenetic reprogramming in migrating primordial germ cells in mice. Development 134, 2627–2638 (2007).

11. S. Guibert, T. Forne, M. Weber, Global profiling of DNA methylation erasure in mouse primordial germ cells. Genome Res 22, 633–641 (2012).

12. S. Gkountela et al., The ontogeny of cKIT+ human primordial germ cells proves to be a resource for human germ line reprogramming, imprint erasure and in vitro differentiation. Nat Cell Biol 15, 113–122 (2013).

13. J. A. Hackett, J. J. Zylicz, M. A. Surani, Parallel mechanisms of epigenetic reprogramming in the germline. Trends Genet 28, 164–174 (2012).

14. S. Seisenberger et al., The dynamics of genome-wide DNA methylation reprogramming in mouse primordial germ cells. Mol Cell 48, 849–862 (2012).

15. S. Saxena, L. Zou, Hallmarks of DNA replication stress. Mol Cell 82, 2298–2314 (2022).

16. L. H. Hartwell, M. B. Kastan, Cell cycle control and cancer. Science 266, 1821–1828 (1994).

17. J. E. Sale, Translesion DNA synthesis and mutagenesis in eukaryotes. Cold Spring Harb Perspect Biol 5, a012708 (2013).

18. J. E. Sale, A. R. Lehmann, R. Woodgate, Y-family DNA polymerases and their role in tolerance of cellular DNA damage. Nat Rev Mol Cell Biol 13, 141–152 (2012).

19. H. Ohmori et al., The Y-family of DNA polymerases. Mol Cell 8, 7–8 (2001).

20. L. Stolk et al., Meta-analyses identify 13 loci associated with age at menopause and highlight DNA repair and immune pathways. Nat Genet 44, 260–268 (2012).

21. F. R. Day et al., Large-scale genomic analyses link reproductive aging to hypothalamic signaling, breast cancer susceptibility and BRCA1-mediated DNA repair. Nat Genet 47, 1294–1303 (2015).

22. K. S. Ruth et al., Genetic insights into biological mechanisms governing human ovarian ageing. Nature 596, 393–397 (2021).

23. R. J. Hill, G. P. Crossan, DNA cross-link repair safeguards genomic stability during premeiotic germ cell development. Nat Genet 51, 1283–1294 (2019).

24. P. Hajkova et al., Genome-wide reprogramming in the mouse germ line entails the base excision repair pathway. Science 329, 78–82 (2010).

25. M. Lutzmann et al., MCM8- and MCM9-deficient mice reveal gametogenesis defects and genome instability due to impaired homologous recombination. Mol Cell 47, 523–534 (2012).

26. K. Sasaki et al., Robust In Vitro Induction of Human Germ Cell Fate from Pluripotent Stem Cells. Cell Stem Cell 17, 178–194 (2015).

27. W. Lin et al., The human REV1 gene codes for a DNA template-dependent dCMP transferase. Nucleic Acids Res 27, 4468–4475 (1999).

28. D. T. Nair, R. E. Johnson, L. Prakash, S. Prakash, A. K. Aggarwal, Rev1 employs a novel mechanism of DNA synthesis using a protein template. Science 309, 2219–2222 (2005).

29. Y. Murakumo et al., Interactions in the error-prone postreplication repair proteins hREV1, hREV3, and hREV7. J Biol Chem 276, 35644–35651 (2001).

30. L. J. Simpson, J. E. Sale, Rev1 is essential for DNA damage tolerance and non-templated immunoglobulin gene mutation in a vertebrate cell line. EMBO J 22, 1654–1664 (2003).

31. J. G. Jansen et al., Strand-biased defect in C/G transversions in hypermutating immunoglobulin genes in Rev1-deficient mice. J Exp Med 203, 319–323 (2006).

32. F. W. Buaas et al., Plzf is required in adult male germ cells for stem cell self-renewal. Nat Genet 36, 647–652 (2004).

33. Y. I. Yeom et al., Germline regulatory element of Oct-4 specific for the totipotent cycle of embryonal cells. Development 122, 881–894 (1996).

34. P. E. Szabo, K. Hubner, H. Scholer, J. R. Mann, Allele-specific expression of imprinted genes in mouse migratory primordial germ cells. Mech Dev 115, 157–160 (2002).

35. A. L. Ross, L. J. Simpson, J. E. Sale, Vertebrate DNA damage tolerance requires the C-terminus but not BRCT or transferase domains of REV1. Nucleic Acids Res 33, 1280–1289 (2005).

36. K. Masuda et al., A critical role for REV1 in regulating the induction of C:G transitions and A:T mutations during Ig gene hypermutation. J Immunol 183, 1846–1850 (2009).

37. E. Ohashi et al., Identification of a novel REV1-interacting motif necessary for DNA polymerase kappa function. Genes Cells 14, 101–111 (2009).

38. M. Khalaj et al., A missense mutation in Rev7 disrupts formation of Polzeta, impairing mouse development and repair of genotoxic agent-induced DNA lesions. J Biol Chem 289, 3811–3824 (2014).

39. M. Pirouz, S. Pilarski, M. Kessel, A critical function of Mad2l2 in primordial germ cell development of mice. PLoS Genet 9, e1003712 (2013).

40. N. Watanabe et al., The REV7 subunit of DNA polymerase zeta is essential for primordial germ cell maintenance in the mouse. J Biol Chem 288, 10459–10471 (2013).

41. S. Shachar et al., Two-polymerase mechanisms dictate error-free and error-prone translesion DNA synthesis in mammals. EMBO J 28, 383–393 (2009).

42. Z. Livneh, O. Ziv, S. Shachar, Multiple two-polymerase mechanisms in mammalian translesion DNA synthesis. Cell Cycle 9, 729–735 (2010).

43. S. Sharma et al., REV1 and polymerase zeta facilitate homologous recombination repair. Nucleic Acids Res 40, 682–691 (2012).

44. A. Bhat, Z. Wu, V. M. Maher, J. J. McCormick, W. Xiao, Rev7/Mad2B plays a critical role in the assembly of a functional mitotic spindle. Cell Cycle 14, 3929–3938 (2015).

45. T. Listovsky, J. E. Sale, Sequestration of CDH1 by MAD2L2 prevents premature APC/C activation prior to anaphase onset. J Cell Biol 203, 87–100 (2013).

46. H. Ghezraoui et al., 53BP1 cooperation with the REV7-shieldin complex underpins DNA structure-specific NHEJ. Nature 560, 122–127 (2018).

47. S. K. Martin, R. D. Wood, DNA polymerase zeta in DNA replication and repair. Nucleic Acids Res 47, 8348–8361 (2019).

48. C. Hoege, B. Pfander, G. L. Moldovan, G. Pyrowolakis, S. Jentsch, RAD6-dependent DNA repair is linked to modification of PCNA by ubiquitin and SUMO. Nature 419, 135–141 (2002).

49. P. Stelter, H. D. Ulrich, Control of spontaneous and damage-induced mutagenesis by SUMO and ubiquitin conjugation. Nature 425, 188–191 (2003).

50. H. Gali et al., Role of SUMO modification of human PCNA at stalled replication fork. Nucleic Acids Res 40, 6049–6059 (2012).

51. M. Mohiuddin et al., SUMOylation of PCNA by PIAS1 and PIAS4 promotes template switch in the chicken and human B cell lines. Proc Natl Acad Sci U S A 115, 12793–12798 (2018).

52. A. Hendel et al., PCNA ubiquitination is important, but not essential for translesion DNA synthesis in mammalian cells. PLoS Genet 7, e1002262 (2011).

53. C. E. Edmunds, L. J. Simpson, J. E. Sale, PCNA ubiquitination and REV1 define temporally distinct mechanisms for controlling translesion synthesis in the avian cell line DT40. Mol Cell 30, 519–529 (2008).

54. P. Langerak, A. O. Nygren, P. H. Krijger, P. C. van den Berk, H. Jacobs, A/T mutagenesis in hypermutated immunoglobulin genes strongly depends on PCNAK164 modification. J Exp Med 204, 1989–1998 (2007).

55. P. Langerak, A. O. Nygren, J. P. Schouten, H. Jacobs, Rapid and quantitative detection of homologous and non-homologous recombination events using three oligonucleotide MLPA. Nucleic Acids Res 33, e188 (2005).

56. S. Roa et al., Ubiquitylated PCNA plays a role in somatic hypermutation and class-switch recombination and is required for meiotic progression. Proc Natl Acad Sci U S A 105, 16248–16253 (2008).

57. B. Payer et al., Generation of stella-GFP transgenic mice: a novel tool to study germ cell development. Genesis 44, 75–83 (2006).

58. B. Pilzecker et al., DNA damage tolerance in hematopoietic stem and progenitor cells in mice. Proc Natl Acad Sci U S A 114, E6875–E6883 (2017).

59. A. Wilson et al., Hematopoietic stem cells reversibly switch from dormancy to self-renewal during homeostasis and repair. Cell 135, 1118–1129 (2008).

60. L. J. Mah, A. El-Osta, T. C. Karagiannis, gammaH2AX: a sensitive molecular marker of DNA damage and repair. Leukemia 24, 679–686 (2010).

61. G. G. Oakley, S. M. Patrick, Replication protein A: directing traffic at the intersection of replication and repair. Front Biosci (Landmark Ed) 15, 883–900 (2010).

62. D. W. Nicholson et al., Identification and inhibition of the ICE/CED-3 protease necessary for mammalian apoptosis. Nature 376, 37–43 (1995).

63. F. Chehrehasa, A. C. Meedeniya, P. Dwyer, G. Abrahamsen, A. Mackay-Sim, EdU, a new thymidine analogue for labelling proliferating cells in the nervous system. J Neurosci Methods 177, 122–130 (2009).

64. M. J. Hendzel et al., Mitosis-specific phosphorylation of histone H3 initiates primarily within pericentromeric heterochromatin during G2 and spreads in an ordered fashion coincident with mitotic chromosome condensation. Chromosoma 106, 348–360 (1997).

65. A. Hagting, M. Jackman, K. Simpson, J. Pines, Translocation of cyclin B1 to the nucleus at prophase requires a phosphorylation-dependent nuclear import signal. Curr Biol 9, 680–689 (1999).

66. J. Pines, T. Hunter, Human cyclins A and B1 are differentially located in the cell and undergo cell cycle-dependent nuclear transport. J Cell Biol 115, 1–17 (1991).

67. A. Gillich et al., Epiblast stem cell-based system reveals reprogramming synergy of germline factors. Cell Stem Cell 10, 425–439 (2012).

68. J. A. Hackett et al., Promoter DNA methylation couples genome-defence mechanisms to epigenetic reprogramming in the mouse germline. Development 139, 3623–3632 (2012).

69. P. W. S. Hill et al., Epigenetic reprogramming enables the transition from primordial germ cell to gonocyte. Nature 555, 392–396 (2018).

70. C. A. Adelman et al., HELQ promotes RAD51 paralogue-dependent repair to avert germ cell loss and tumorigenesis. Nature 502, 381–384 (2013).

71. K. Nishimura et al., Mcm8 and Mcm9 form a complex that functions in homologous recombination repair induced by DNA interstrand crosslinks. Mol Cell 47, 511–522 (2012).

72. A. I. Agoulnik et al., A novel gene, Pog, is necessary for primordial germ cell proliferation in the mouse and underlies the germ cell deficient mutation, gcd. Hum Mol Genet 11, 3047–3053 (2002).

