## Supplementary Material for "Primordial germ cell DNA demethylation and development require DNA translesion synthesis"

### Supplementary Figures:

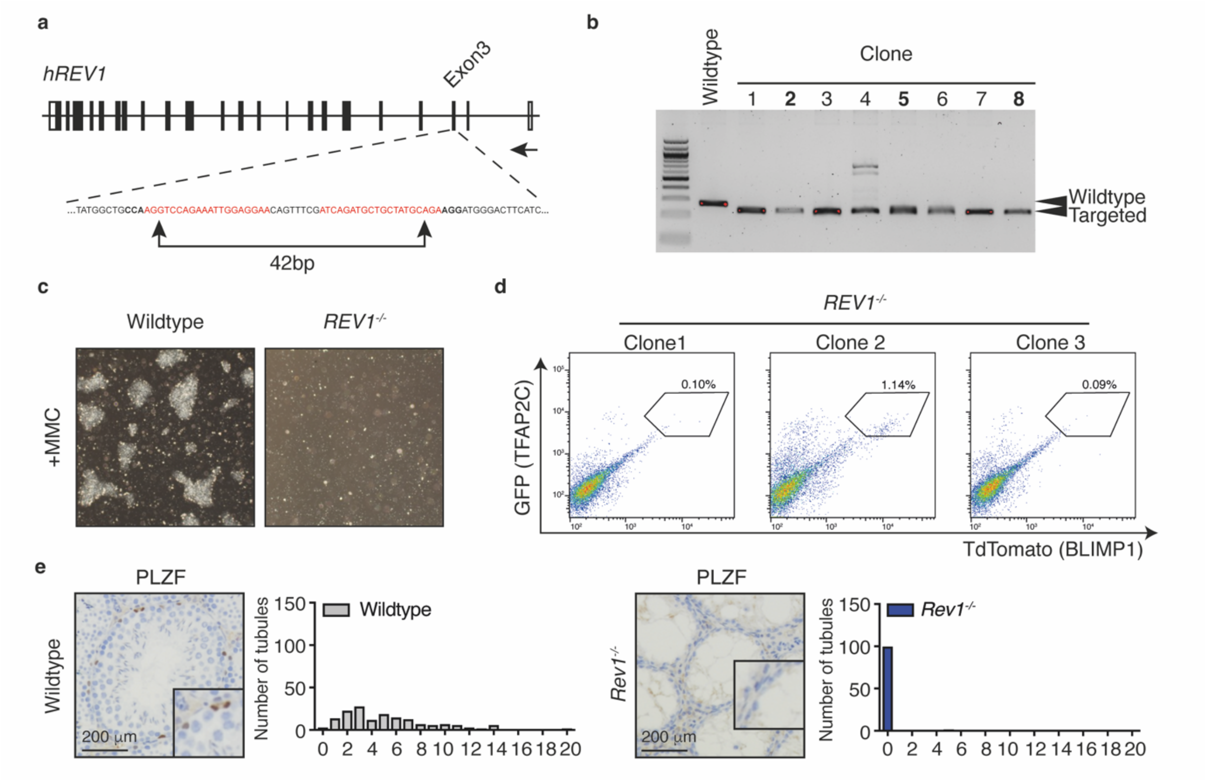


**Supplementary Figure 1. Generation of REV1-deficient hiPSCs and characterization of *Rev1$^{-/-}$* mouse testes.**

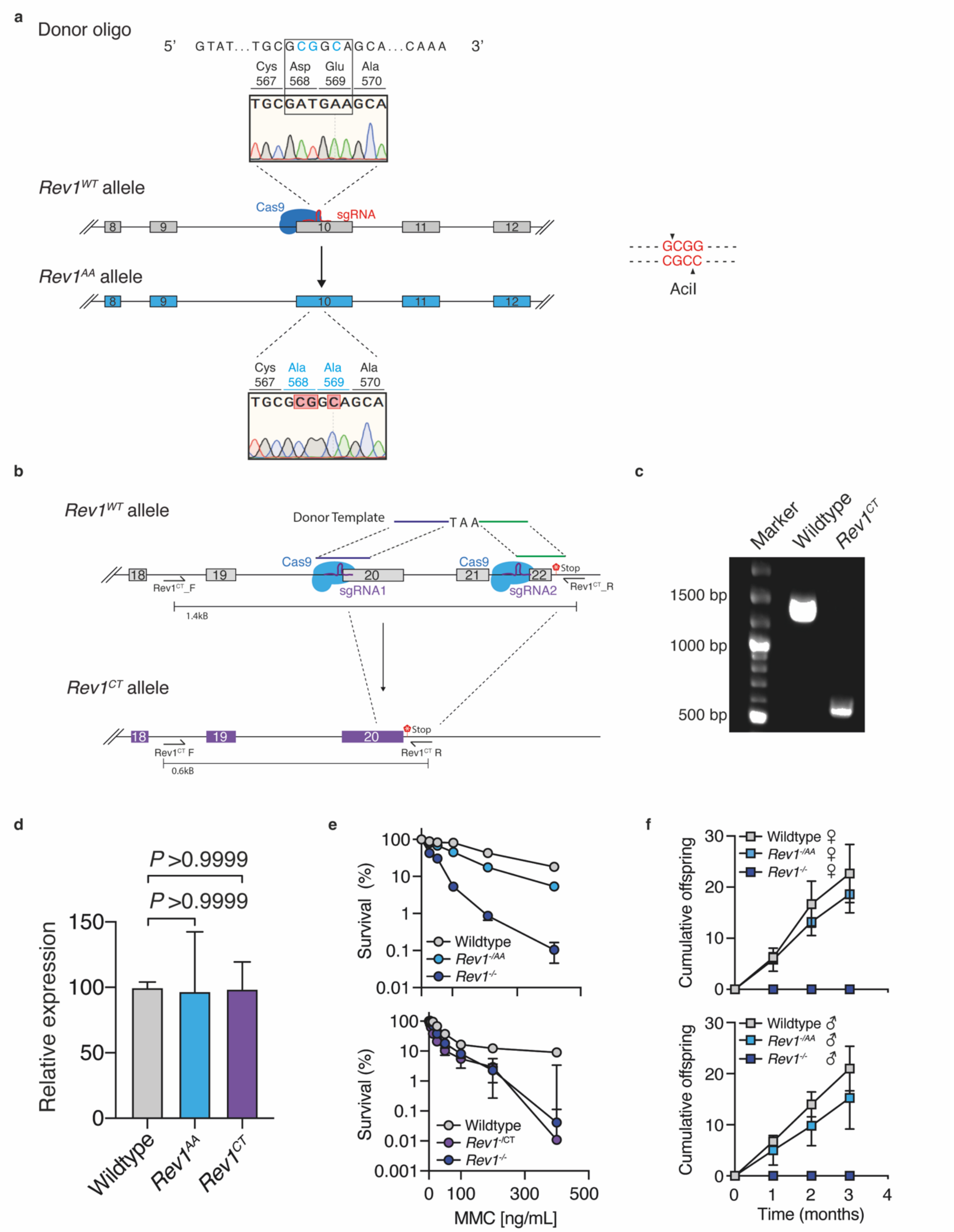


**Supplementary Figure 2. Generation and validation of Rev1AA and Rev1CT alleles by CRISPR-Cas9 genome editing.**

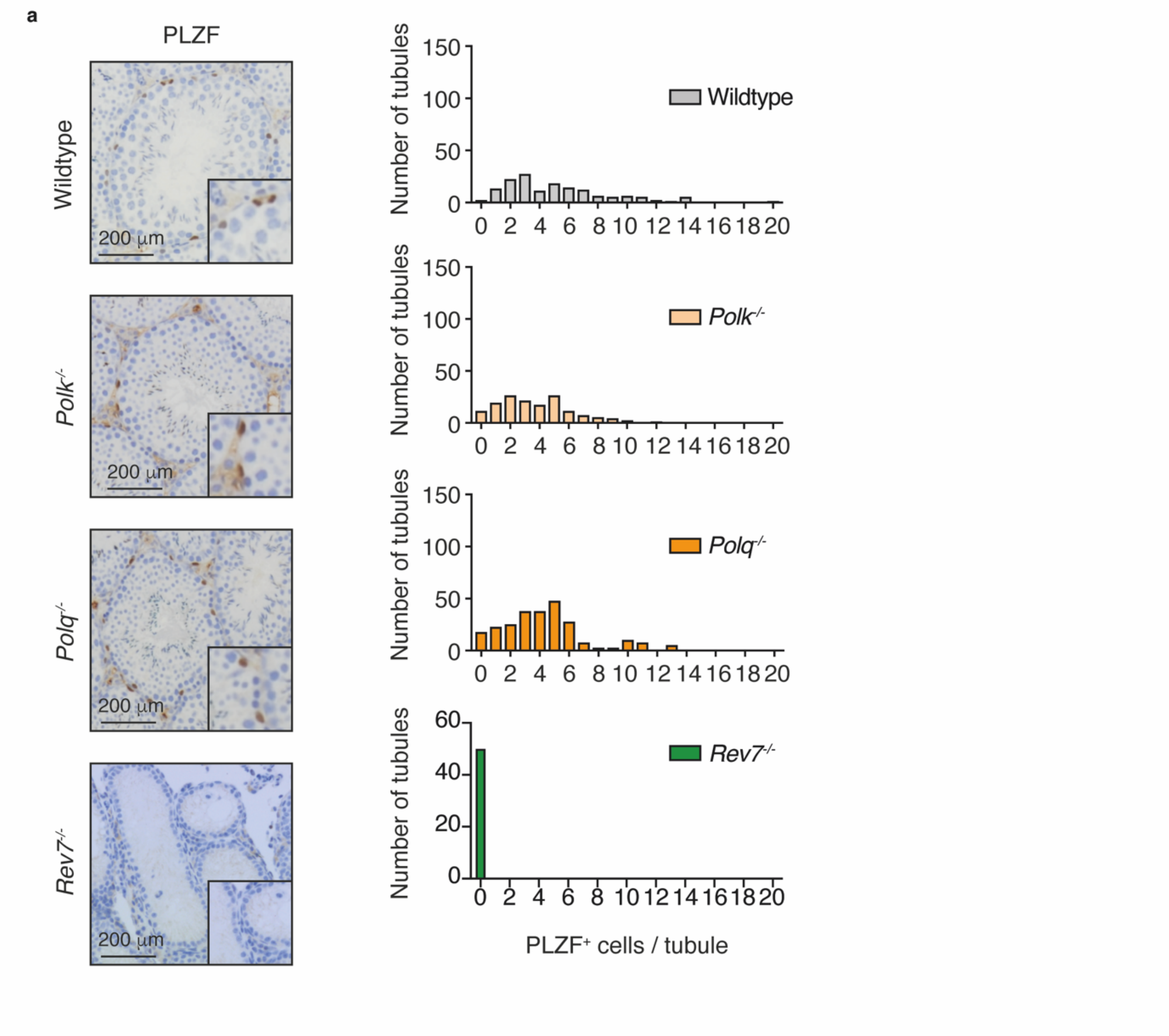


20 **Supplementary Figure 3. Lack of PLZF$^{+}$ cells in the testes of REV7-deficient mice.**

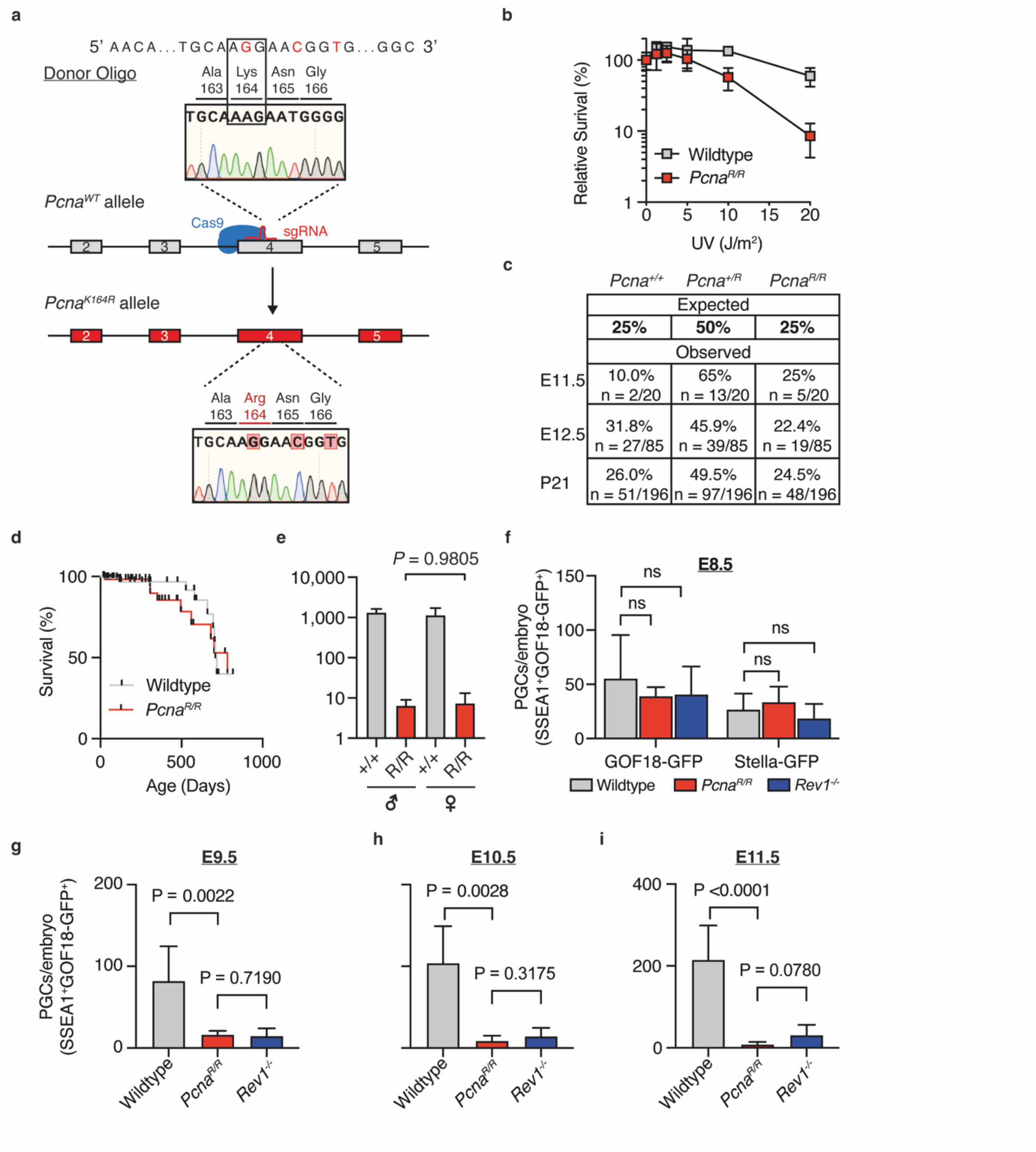


**Supplementary Figure 4. Generation and characterisation of *Pcna*$^{R/R}$ mice.**

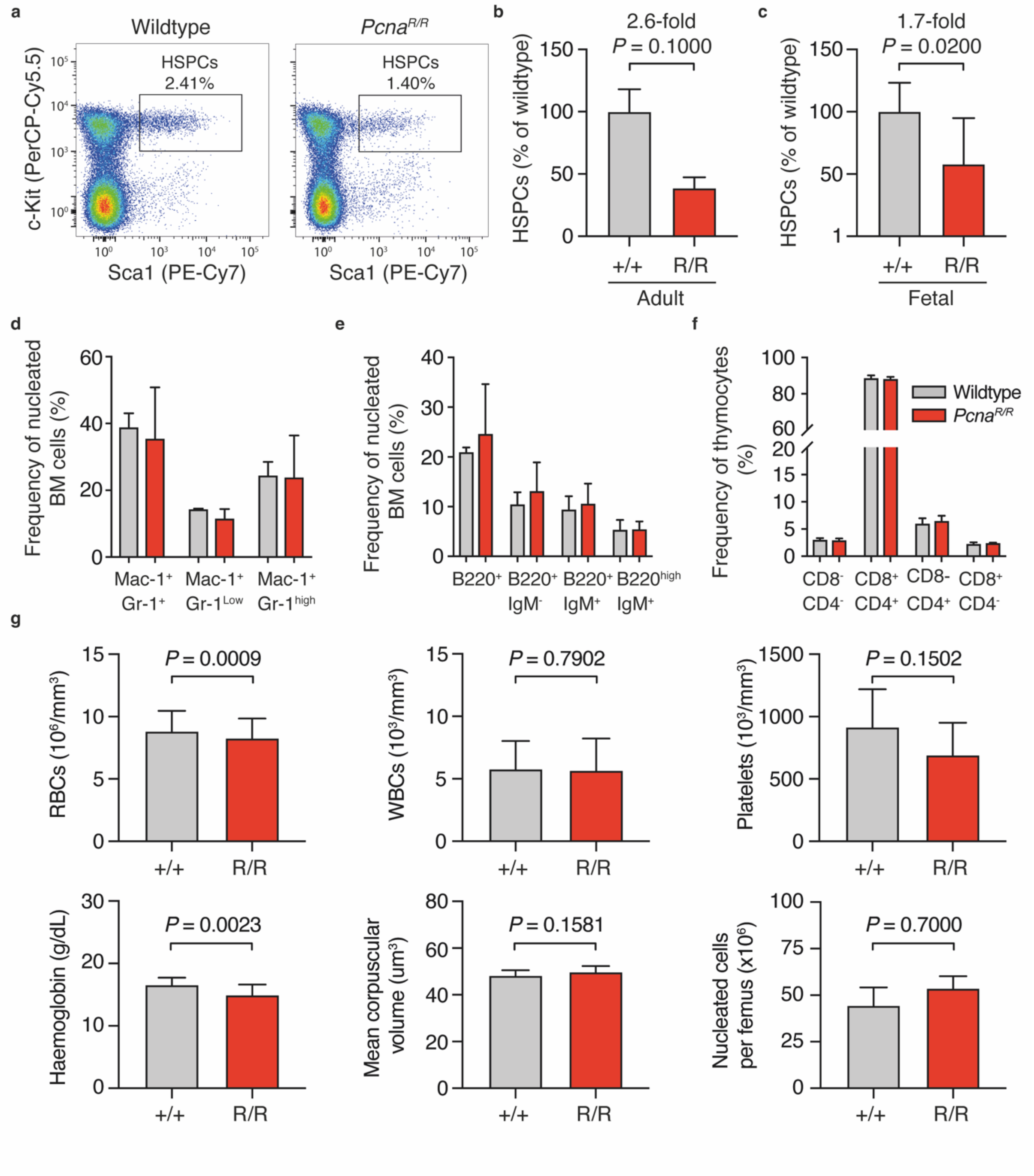


**Supplementary Figure 5. Normal haematopoiesis in *Pcna*$^{R/R}$ mice.**

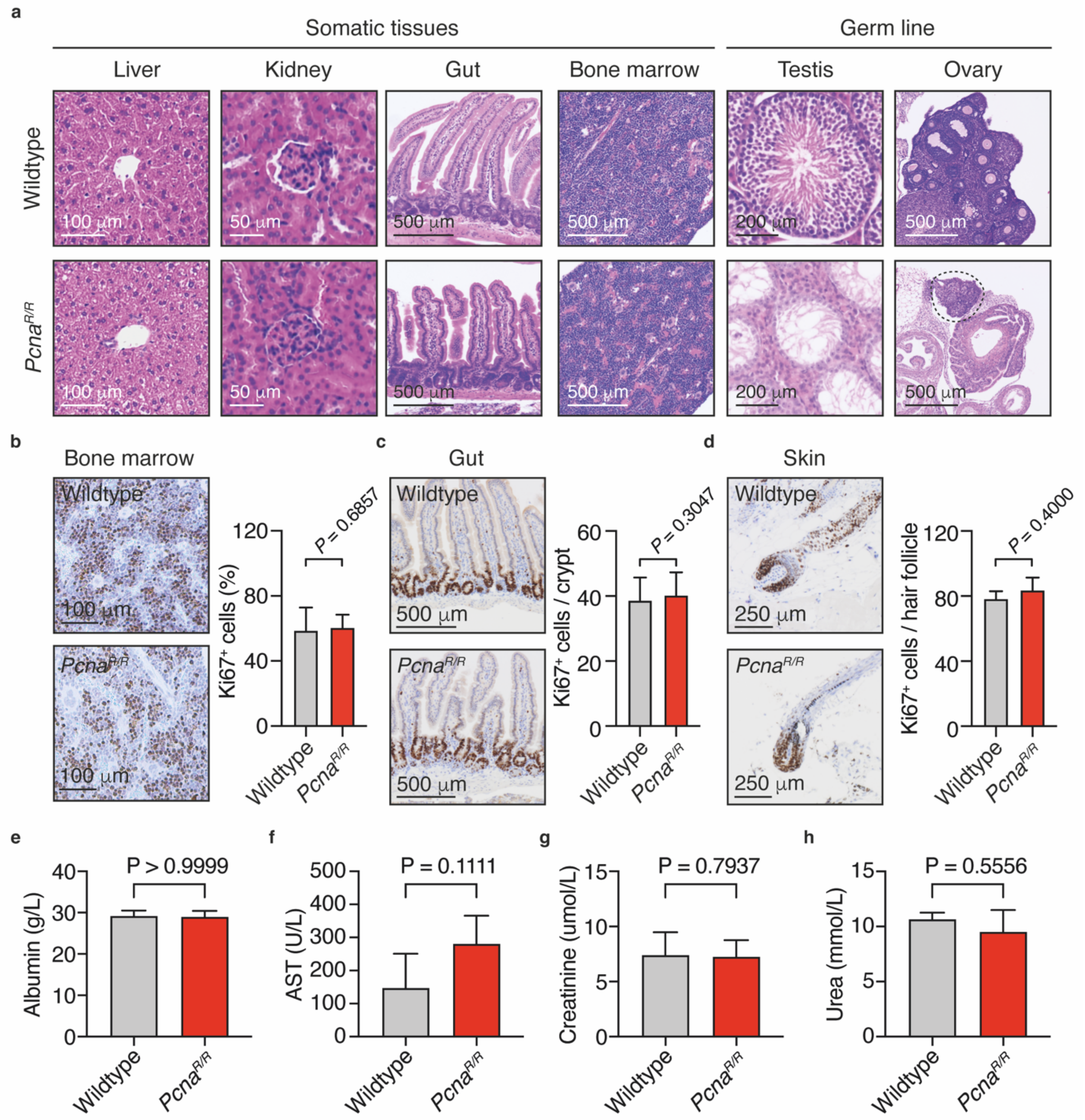


**Supplementary Figure 6. Analysis of somatic tissues from *Pcna*$^{R/R}$ mice.**

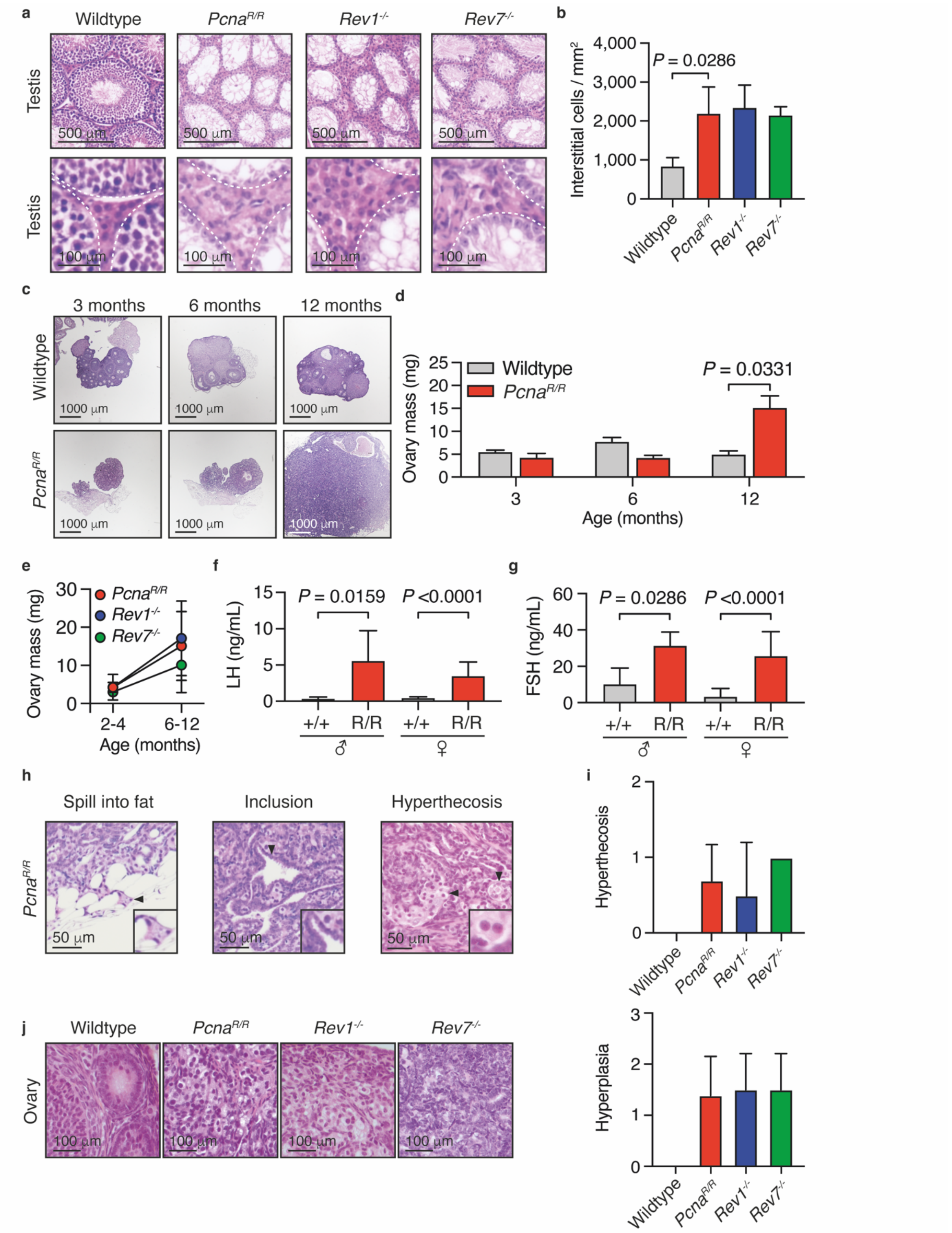


**Supplementary Figure 7. The hypothalamic-pituitary-gonadotrophic axis is disrupted in TLS-deficient adults.**

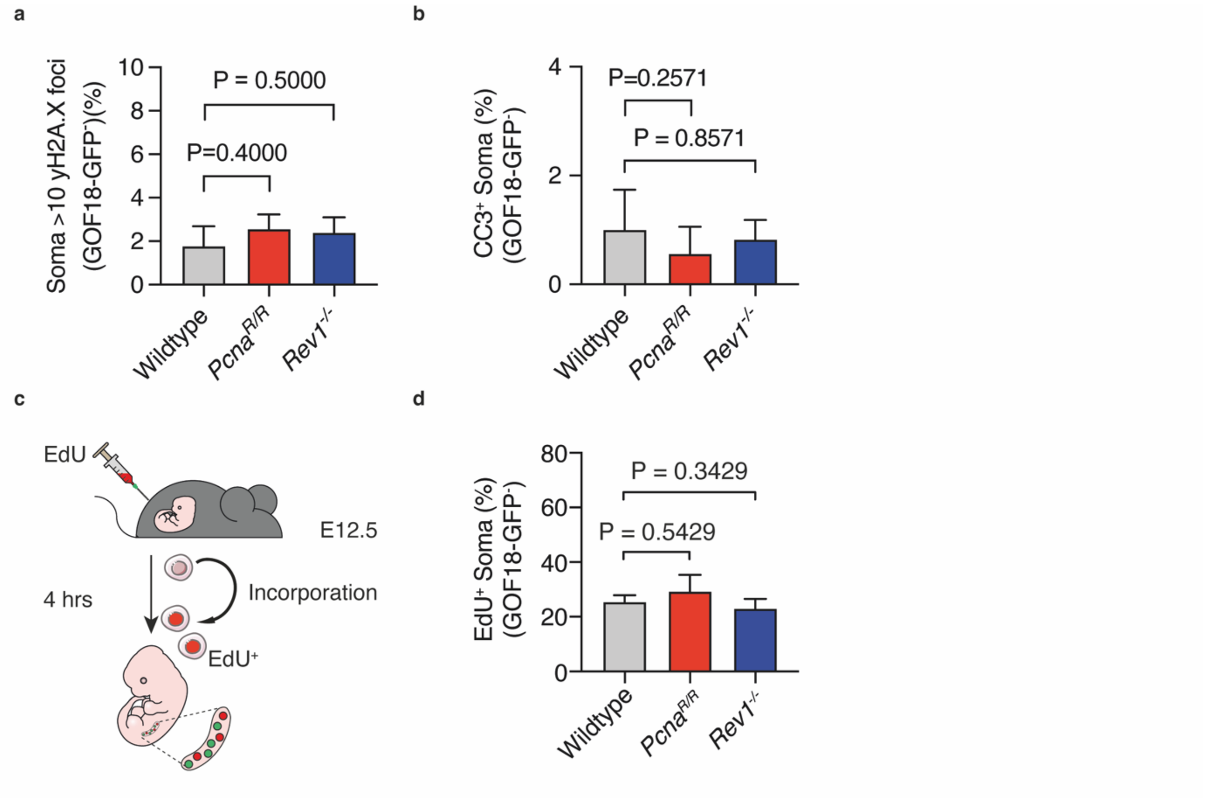


**Supplementary Figure 8. PCNA K164 and REV1 are dispensable in gonadal somatic cells.**

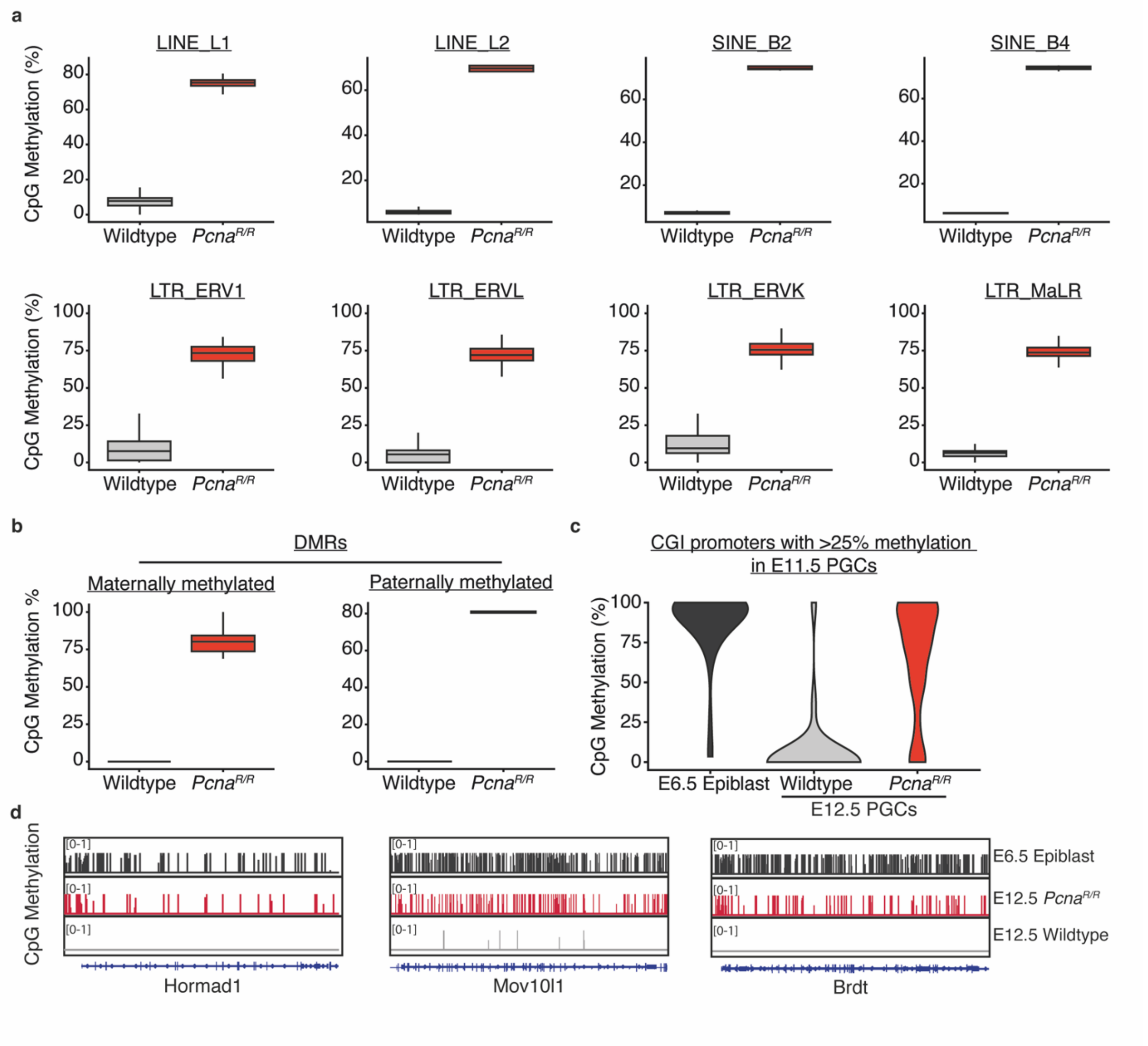


**Supplementary Figure 9. Global retention of methylation in E12.5 *Pcna*$^{R/R}$ PGCs.**

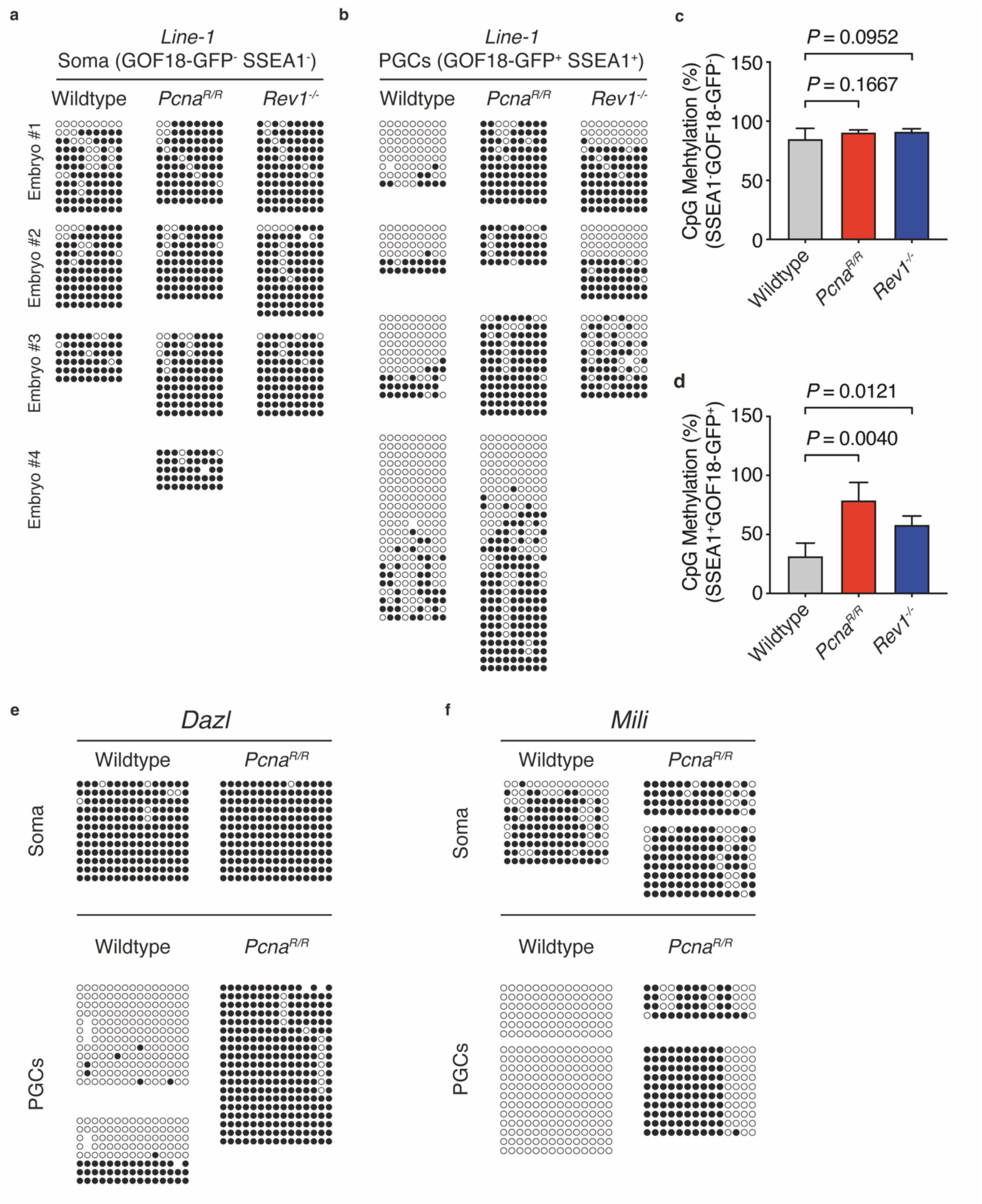


35 **Supplementary Figure 10.** ***Pcna**$^{R/R}$* **and** ***Rev1**$^{-/-}$* **PGCs retain DNA methylation.**

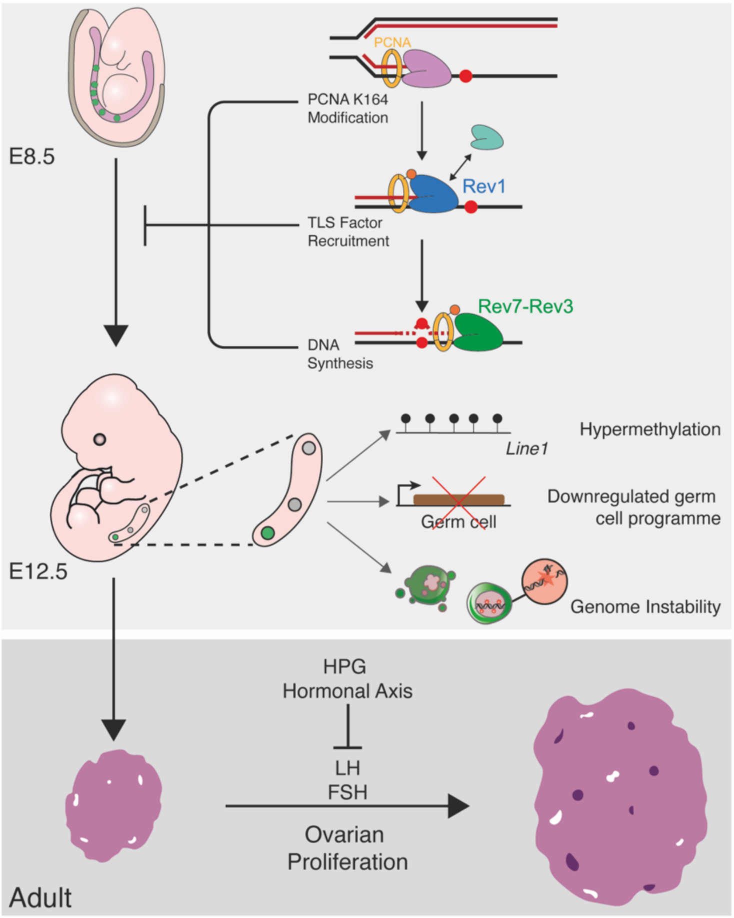


**Supplementary Figure 11. DNA translesion synthesis is essential for PGC proliferation, DNA demethylation, and fertility.**

**Supplementary Figure 1. Generation of REV1-deficient hiPSCs and characterization of *Rev1*$^{-/-}$ mouse testes**

**(a)** Schematic representation of the *REV1* locus on chromosome 2 and the two gRNA targeting sites.

**(b)** Gel image of *REV1* PCR amplicons, using gDNA isolated from single cell derived colonies after electroporation. Clones analyzed are numerated from 1 to 8. Size of the wildtype amplicon is 281bp and of the targeted amplicon is 239bp. Clone 4 contains bands of unexpected size, and was therefore not used in further analysis.

**(c)** Representative brightfield images of wildtype and *REV1*$^{-/-}$ hiPSCs treated with 5 ng/ml mitomycin c (MMC) for 48 hours.

**(d)** Representative flow cytometry plots of *TFAPC*-GFP and *BLIMP1*-tdTomato positive hPGCLCs at day 4 of aggregate differentiation from from *REV1*$^{-/-}$ clones.

**(e)** Right: PLZF-stained testes from 8–12-week-old wildtype and *Rev1*$^{-/-}$ mice (similar results were obtained from 3 independent animals per genotype and sex). Left: Frequency of PLZF$^{+}$ cells per seminiferous tubule of 8-12-week-old mice (the data shown represent the median and interquartile range; n=150 tubules per genotype, 50 per genotype).

**Supplementary Figure 2. Generation and validation of Rev1AA and Rev1CT alleles by CRISPR-Cas9 genome editing**

**(a)** Generation of the *Rev1*$^{AA}$ allele in mouse embryonic stem cells (mESCs). Sanger sequencing profiles of the wildtype (top) and targeted *Rev1*$^{AA}$ (bottom) locus from confirming successful mutagenesis of the catalytic residues D568 and E569 to alanines (D568A and E569A).

**(b)** Generation of the *Rev1*$^{CT}$ allele by CRISPR-Cas9 genome editing in mouse zygotes. Two sgRNAs (sgRNA1 and sgRNA2) were used to target Cas9 activity to sites either side of REV1's C-terminus. A template DNA molecule (donor template) was then used to insert a stop codon (TAA) and truncate the C-terminus.

**(c)** Verification of gene targeting by long-range PCR. Oligonucleotide pairs were designed either side of the DNA encoding the C-terminal 100 amino acids of REV1. In the wildtype *Rev1* allele a 1,500 bp band is amplified. This is truncated to a 600 bp product in the *Rev1*$^{CT}$ allele.

**(d)** RT-qPCR of the *Rev1* transcript in wildtype, *Rev1*$^{-/AA}$ and *Rev1*$^{-/CT}$ mouse embryonic fibroblasts (n=3 independent experiments).

**(e)** Sensitivity of wildtype, *Rev1*$^{-/-}$ and *Rev1*$^{-/AA}$ (top) or *Rev1*$^{-/CT}$ (bottom) cells to MMC (data represent 3 independent experiments each carried out in triplicate).

**(e)** Cumulative number of offspring when mutant female (top) or male (bottom) mice were mated with wildtype mates and checked for evidence of copulation (n=3 independent mice per genotype and sex).

**Supplementary Figure 3. Lack of PLZF$^{+}$ cells in the testes of REV7-deficient mice**

**(a)** Representative images of testis sections stained for PLZF and the quantification of PLZF$^{+}$ cells per seminiferous tubule of wildtype, *Polk*$^{-/-}$, *Polq*$^{-/-}$ and *Rev7*$^{-/-}$ mice (the data shown represent the median and interquartile range; n=150 tubules per genotype, 50 per animal).

**Supplementary Figure 4. Generation and characterisation of $Pcna^{R/R}$ mice**

**(a)** Generation of the $Pcna^{K164R}$ mutation by CRISPR-Cas9 genome editing in mouse zygotes. Sanger sequencing profiles of the wildtype (top) and targeted (bottom) *Pcna* locus confirming successful mutagenesis.

**(b)** Sensitivity of wildtype and $Pcna^{R/R}$ MEFs to ultraviolet (*57*) irradiation (data represent 3 independent experiments each carried out in triplicate).

**(c)** $Pcna^{R/R}$ embryos are observed at the expected Mendelian ratios throughout development (E11.5 and E12.5) and at 21 days postpartum (P21).

**(d)** Kaplan-Meier survival curve for a cohort of wildtype and $Pcna^{R/R}$ mice (n=125 wildtype mice, 62 $Pcna^{R/R}$ mice).

**(e)** Quantification of PGCs by flow cytometry from male and female embryos at E12.5 (*P* value calculated by two-tailed Mann-Whitney *U*-test, the data shown represent the mean and s.d.; *n*=13, 17, 7 and 10 independent embryos, left to right).

**(f)** Quantification of PGCs by flow cytometry from wildtype, $Rev1^{-/-}$ and $Pcna^{R/R}$ embryos at E8.5 (wildtype, n = 18 and 6; $Rev1^{-/-}$, n = 5 and 2; $Pcna^{R/R}$, n = 2 and 4, independent embryos, left to right).

**(g-i)** Quantification of PGCs by flow cytometry through development (E9.5-11.5) from wildtype, $Pcna^{R/R}$ and $Rev1^{-/-}$ embryos (E9.5, n=12, 4 and 6; E10.5, n=9, 4 and 5; E11.5, n=7, 16 and 4; left to right).

**Supplementary Figure 5. Normal haematopoiesis in *Pcna*$^{R/R}$ mice**

**(a)** Representative flow cytometry plots of hematopoietic stem and progenitor cells (HSPCs) of wildtype and *Pcna*$^{R/R}$ mice.

**(b)** Quantification of HSPCs (Lin-Kit$^{+}$Sca-1$^{+}$) by flow cytometry from 8-12-week-old wildtype and *Pcna*$^{R/R}$ mice (n=3 independent mice per genotype).

**(c)** Quantification HSPCs (Lin-Kit$^{+}$Sca-1$^{+}$) by flow cytometry from wildtype and *Pcna*$^{R/R}$ embryos at E12.5 (n=4 independent mice per genotype).

**(d)** Quantification of myeloid cell by flow cytometry from bone marrow of 8-12-week-old wildtype and *Pcna*$^{R/R}$ mice stained for the myeloid markers Mac-1 and Gr-1 and the quantification of distinct populations (n=3 independent mice per genotype).

**(e)** Quantification of B-cell maturation by flow cytometry from bone marrow of 8-12-week-old wildtype and *Pcna*$^{R/R}$ mice stained for the B cell markers B220 and IgM and the quantification of distinct populations (n=3 independent mice per genotype).

**(f)** Quantification of T-cell development by flow cytometry from the thymus of 8-12-week-old wildtype and *Pcna*$^{R/R}$ mice stained for the T cell markers CD4 and CD8 and the quantification of distinct populations (n=3 independent mice per genotype).

**(g)** Whole blood counts and quantification of nucleated cells per femurs of 8-12-week-old wildtype and *Pcna*$^{R/R}$ mice (n=18 wildtype and 21 *Pcna*$^{R/R}$ mice; RBC, red blood cells; WBC, white blood cells).

**Supplementary Figure 6. Analysis of somatic tissues from *Pcna*$^{R/R}$ mice**

**(a)** H&E-stained sections of liver, kidney, gut and bone marrow sections (left) and testis and ovary sections (right) from wildtype and *Pcna*$^{R/R}$ adult mice (similar observations were made in 3 independent animals per genotype).

**(b)** Representative images of bone marrow sections from 8-12-week-old wildtype and *Pcna*$^{R/R}$ mice stained for Ki67 and quantification of the frequency of Ki67$^{+}$ cells (n=3 independent mice per genotype).

**(c)** Representative images of ileum sections stained for Ki67 from 8-12-week-old wildtype and *Pcna*$^{R/R}$ mice and quantification of Ki67$^{+}$ cells per crypt of the ileum n=3 independent mice per genotype).

**(d)** Representative images of skin sections from 8-12-week-old wildtype and *Pcna*$^{R/R}$ mice stained for Ki67 and quantification of Ki67$^{+}$ cells per hair follicle bulge (n=3 independent mice per genotype).

**(E-H)** Serum levels of albumin, AST, creatinine and urea (wildtype n=5;*Pcna*$^{R/R}$ n=4).

**Supplementary Figure 7. The hypothalamic-pituitary-gonadotrophic axis is disrupted in TLS-deficient adults**

**(a)** Representative images of H&E-stained testis sections from 6-12-month-old wildtype, *Pcna*$^{R/R}$, *Rev1*$^{-/-}$ and *Rev7*$^{-/-}$ mice

**(b)** Quantification of interstitial cells, mostly Leydig cells (n=3 independent mice per genotype).

**(c)** Representative H&E-stained sections of ovaries from 3-, 6- and 12-month-old wildtype and *Pcna*$^{R/R}$ mice (similar results were obtained from 3 independent animals per genotype).

**(d)** Quantification of ovary mass from 3-, 6- and 12-month-old wildtype and *Pcna*$^{R/R}$ mice ( n=18, 2, 10, 5, 6, and 13 independent animals, left to right).

**(e)** Quantification of ovarian mass in 2-4- and 6–12-month-old *Pcna*$^{R/R}$, *Rev1*$^{-/-}$ and *Rev7*$^{-/-}$ mice (2-4 months, n=3, 2 and 4, 6-12 months, n=12, 2 and 2 independent animals for *Pcna*$^{R/R}$, *Rev1*$^{-/-}$ and *Rev7*$^{-/-}$ respectively).

**(f)** Quantification of serum luteinizing hormone (*46*) concentrations in male and female wildtype and *Pcna*$^{R/R}$mice (n=5, 4, 14 and 13 independent animals, left to right).

**(g)** Quantification of serum follicle stimulating hormone (FSH) concentration in male and female wildtype and *Pcna*$^{R/R}$ mice (n=4, 4, 14 and 13 independent animals, left to right).

**(h)** Representative H&E-stained ovaries from 12-month-old *Pcna*$^{R/R}$ mice demonstrating pathological changes of ovarian stromal hyperplasia, including overspill of hyperplastic stromal cells into the adjacent fat at the hilum (left), surface epithelial inclusions (center) and areas of hyperthecosis (right).

**(i)** Pathological assessment of stromal cell hyperthecosis of ovaries from 6-12-month-old wildtype, *Pcna*$^{R/R}$, *Rev1*$^{-/-}$ and *Rev7*$^{-/-}$ mice (n=14, 13, 2 and 2 independent animals, left to right).

**(j)** Representative images of H&E-stained ovary sections from 6-12-month-old wildtype, *Pcna*$^{R/R}$, *Rev1*$^{-/-}$ and *Rev7*$^{-/-}$ mice and the assessment of stromal hyperplasia (n=3 independent mice per genotype).

**Supplementary Figure 8. PCNA K164 and REV1 are dispensable in gonadal somatic cells**

**(a)** Frequency of somatic cells ($GFP^-$) with >10 ɣ-H2A.X foci per nucleus from wildtype, $Pcna^{R/R}$ and $Rev1^{-/-}$ embryos at E12.5 (n=3 for all genotypes).
**(b)** Quantification of the frequency of cleaved-caspase 3 (CC3)-positive somatic cells from wildtype, $Pcna^{R/R}$ and $Rev1^{-/-}$ embryos at E12.5 (n=4, 3 and 3, left to right).
**(c)** Schematic of EdU pulse experiment. Pregnant dams were injected with EdU and culled 4 hrs later to harvest E12.5 embryos. The genital ridges of these embryos were analyzed for EdU incorporation using IF.
**(d)** Quantification of the frequency of $EdU^+$ somatic cells of the genital ridge from wildtype, $Pcna^{R/R}$ and $Rev1^{-/-}$ embryos at E12.5 following a 4-hour EdU pulse (n=3 per genotype).

**Supplementary Figure 9. Global retention of methylation in E12.5 *Pcna*$^{R/R}$ PGCs**

**(a)** Quantification of DNA CpG methylation across repeat elements in the genomes of wildtype E6.5 epiblast cells and E12.5 wildtype and *Pcna*$^{R/R}$ PGCs. The percentage methylation is calculated individual locus.

**(b)** Quantification of DNA CpG methylation across maternally and paternally demethylated DMRs in the genomes of wildtype E6.5 epiblast cells and E12.5 wildtype and *Pcna*$^{R/R}$ PGCs. The percentage methylation is calculated over individual DMRs.

**(c)** Quantification of DNA CpG methylation in wildtype E6.5 epiblast cells and E12.5 wildtype PGCs and *Pcna*$^{R/R}$ PGCs across CGI-containing promoters which show >25% methylation at E11.5 as identified previously (*14*).

**(d)** IGV visualization of CpG methylation across selected GRR genes in E6.5 epiblast cells from wildtype embryos, E12.5 wildtype PGCs and E12.5 *Pcna*$^{R/R}$ PGCs. The plots represent the distribution of CpG methylation across genes segmented in 0.1 Kbp genomic windows.

**Supplementary Figure 10. *Pcna*$^{R/R}$ and *Rev1*$^{-/-}$ PGCs retain DNA methylation**

**(a)** Genomic bisulfite sequencing reads of the CpG-rich region of the *Line-1* element from FACS-purified somatic cells from E12.5 wildtype, *Pcna*$^{R/R}$ and *Rev1*$^{-/-}$ embryos (filled = methylated CpG, open = unmethylated CpG).

**(b)** Genomic bisulfite sequencing reads of the CpG-rich region of the *Line-1* element from FACS-purified PGCs from E12.5 wildtype, *Pcna*$^{R/R}$ and *Rev1*$^{-/-}$ embryos

**(c)** Quantification of methylated CpG dinucleotides in the *Line-1* element in somatic cells from E12.5 wildtype, *Pcna*$^{R/R}$ and *Rev1*$^{-/-}$ embryos (n as shown in (a)).

**(d)** Quantification of methylated CpG dinucleotides in the *Line-1* element in PGCs from E12.5 wildtype, *Pcna*$^{R/R}$ and *Rev1*$^{-/-}$ embryos (n as shown in (b)).

**(e)** Genomic bisulfite sequencing reads of the CpG-rich region of the *Dazl* gene promoter element from FACS-purified PGCs from E12.5 wildtype and *Pcna*$^{R/R}$ embryos.

**(f)** Genomic bisulfite sequencing reads of the CpG-rich region of the *Mili* gene promoter element from FACS-purified PGCs from E12.5 wildtype and *Pcna*$^{R/R}$ embryos.

**Supplementary Figure 11. DNA translesion synthesis is essential for PGC proliferation, DNA demethylation, and fertility**

Model for the requirement of TLS factors in mammalian embryonic germ cell development. In the absence of core TLS factors, PGCs display a failure in key aspects of the embryonic germ cell program. This leads to an absence of mature gametes in adulthood which disrupts hormonal signaling. (HPG: Hypothalamic–pituitary–gonadal axis, LH: luteinizing hormone,FSH: follicle stimulating hormone)

### Supplementary Tables:

215

| Supp. Table 1 | Genotyping |
|---|---|
| PcnaK164R | Custom Taqman |
| Rev1AA | Taqman |
| GOF18-GFP & Stella-GFP-1 | AGTGCTTCAGCCGCTACC |
| GOF18-GFP & Stella-GFP-2 | GAAGATGGTGCGCTCCTG |
| GOF18-GFP & Stella-GFP-Pr | FAM-TTCAAGTCCGCCATGCCCGAA-TAMRA |
| Rev1_1 | ATTGTGAGTCTCTAGCGTTTG |
| Rev1_2 | GCTGGAATTGAAATTCTAGG |
| Rev1_3 | GCTTCCATTGCTCAGCGGTG |
| Rev1CT_1 | GTTGTACAGCTGAGCTCGGA |
| Rev1CT_2 | TACCTCACAAGCACTGCTGG |
| Rev1CT_3 | AGCAGTCGGTGGAGTCTGTA |
| PolQ_1 | TGCAGTGTACAGATGTTACTTTT |
| PolQ_2 | TGGAGGTAGCATTTCTTCTC |
| PolQ_3 | TCACTAGGTTGGGGTTCTC |
| PolQ_4 | CATCAGAAGCTGACTCTAGAG |

| | |
|---|---|
| PolK_1 | CTGATGTGACCGCTGTTAAATGTTG |
| PolK_2 | CTGTGGAGATGCCTTAGCGG |
| PolK_3 | GATCCTGCAATCAATAGCTCACGG |
| Rev7_1 | TCCAGGACACACTCCACTGC |
| Rev7_2 | CGTTCTGCAAGCACAGGAAC |
| Rev7_3 | TCGTGGTATCGTTATGCGCC |

**Supplementary Table 1. Assays used to genotype mice.**

| Supp. Table 2 | CRISPR |
|---|---|
| hREV1 gRNA1 | TTCCTCCAATTTCTGGACCT |
| hREV1 gRNA2 | ATCAGATGCTGCTATGCAGA |
| hREV1-screen-F | TGGTCACTAGCACAGAATAAGGT |
| hREV1-screen-R | ACATGCCAAAATAGGGTTAAGTAAA |
| Pcna Donor | GAACAGGAGTACAGCTGTGTAATAAAGATGCCGTCGGGTGAATTTGCACGTATATGCCGAGACCTTAGCCACATTGGAGATGCTGTTGTGATATCCTGTGCAAGGAACGGTGTGAAGTTTTCTGCAAGTGGAGAGCTTGGCAATGGGAACATTAAGTTGTCACAAACAAGTAATGTGGATAAAGAAGAGGAGGCG |
| Pcna sgRNA | TGATATCCTGTGCAAAGAAT |
| Rev1AA Donor | TTTGGTCTTGTCTTTGATTTCAATACGGAGGGCAGCTGCAAACTCCTCAGGAGAAAGTTTCGTCTCTGCAAGGATGTCCGTGACGTCAATCAGTGCTGCCGCGCAGCTGACAGCTTCGATGCTGTGTGTGTAGCTATCAGGACAAAGAGAAGGCAGTGAGTGAGTCACACCTAGGGGCGGCTCTGGGAGTTGTCAGATAC |
| Rev1AA sgRNA | CCTCGATGCTGTGTGTGTAGCTA |
| Rev1CT Donor | CAAAATACTGACTGTTACTAGATTTTAATGTACTTGACACTGTCTGAATACAGCATTCTCGTTCACTTCCTCAACAGGAAGAAGTCTCGGCTTCTACCCTAATGGTTTTACAGCAGACTTATGGAAGCACACTGAAAGTGACCTGACTGCTGTGCAGAGGGCCTGGGGCTCTCTGCGCTGTGCCAGCAGTGCTTGTGAGG |
| Rev1CT sgRNA1 | AATACGACCAGAATGGATTG |
| Rev1CT sgRNA2 | TAAGGCACCATTTGAACTAT |
| Rev1CT-Seq-F | GTTGTACAGCTGAGCTCGGA |
| Rev1CT-Seq-R | TACCTCACAAGCACTGCTGG |

**Supplementary Table 2: Oligonucleotide sequences for CRISPR editing and validation.**

| Supp. Table 3 | qPCR |
|---|---|
| Gapdh | NM_008084.2 |
| Ddx4 | Mm00802445_m1 |
| Nanos3 | Mm00808138_m1 |
| Prdm1 | Mm01187285_m1 |
| Polq | Mm00712819_m1 |
| Rev1 | Mm0045983_m1 |
| Polk | Mm01282564_m1 |
| Rev7 | Mm00510936_m1 |
| Stella_F | GCTAACCCTAAACCCCGGTGT |
| Stella_R | CAATGCGGTTCCGTAGACTGC |
| Fragilis_F | CAGCACCTTGGTCCTCAGCA |
| Fragilis_R | CAGGACCGGAAGTCGGAATC |
| Dazl_F | TCTTTGCCAGATATGGCTCAGT |
| Dazl_R | CTTCTGCACATCCACGTCATTA |
| Mili_F | TTGGCCTCAAGCTCCTAGAC |
| Mili_R | GAACATGGACACCAAACCTACA |

| | |
|---|---|
| Sycp3_F | GCAGTCTAGAATTGTTCAGAGCCAGA |
| Sycp3_R | TCCAAACTCTTTATGAACTGCTCGTG |
| Mael_F | TGGCCACTCTCTTTGGAATC |
| Mael_R | GCATTTCCAATTCTTCCAGC |
| Mov10l1_F | CGCTGTGACGAGTACAGTG |
| Mov10l1_R | CTGACAACCCTTTGCTAGAGTTT |
| Hormad1_F | GGCTCCTAGCTGTTTCAGTATCT |
| Hormad1_R | TTGTCCCATAAGCACGTTCTG |
| Brdt_F | AGTGGGCGGTTGACGAATC |
| Brdt_R | AGTCAGGCAGCTTTAGTTTCAC |
| Tet1_F | GAGCCTGTTCCTCGATGTGG |
| Tet1_R | CAAACCCACCTGAGGCTGTT |
| Dnmt1_F | AAGAATGGTGTTGTCTACCGAC |
| Dnmt1_R | CATCCAGGTTGCTCCCCTTG |

**Supplementary Table 3: Assays used for gene expression analyses.**
